## Supplementary Results for "Newly trained navigation and verbal memory skills elicit changes in task-related networks but not brain structure"

**Supplementary Information**

This supplement includes:

Supplementary Figures 1-8.

Supplementary Tables 1-16.

Supplementary Notes 1-7.

**Supplementary Figures:**

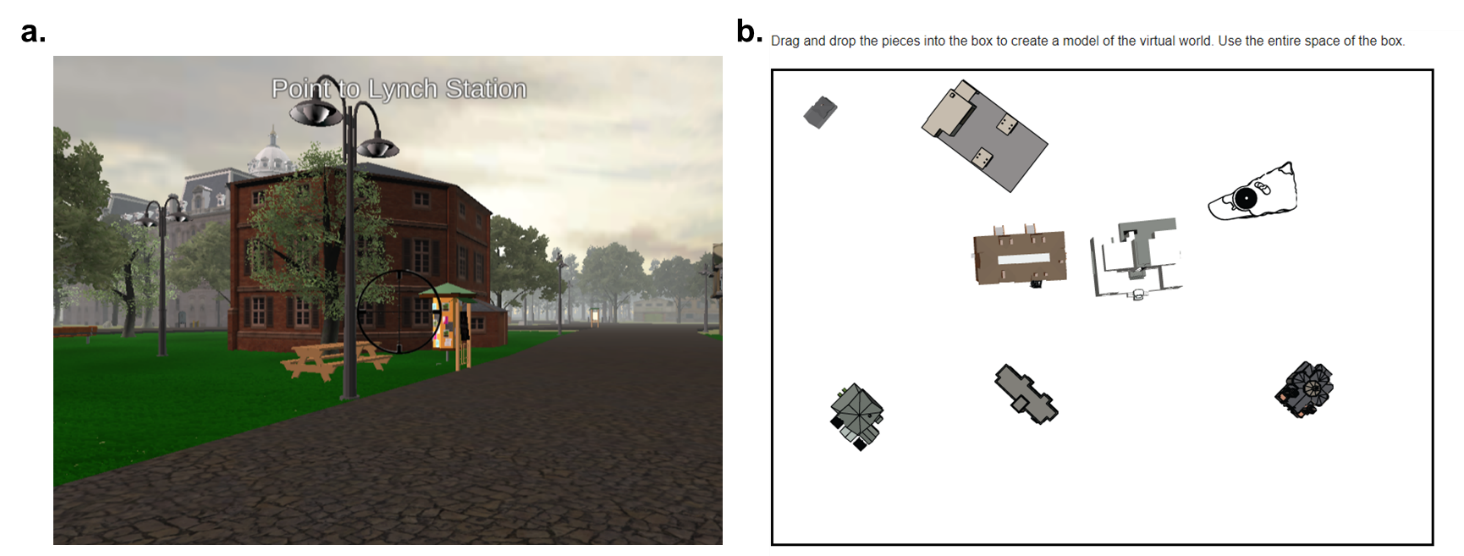

Supplementary Figure 1. Schematic representation of the Navigation Pointing task and Navigation Model Building task.

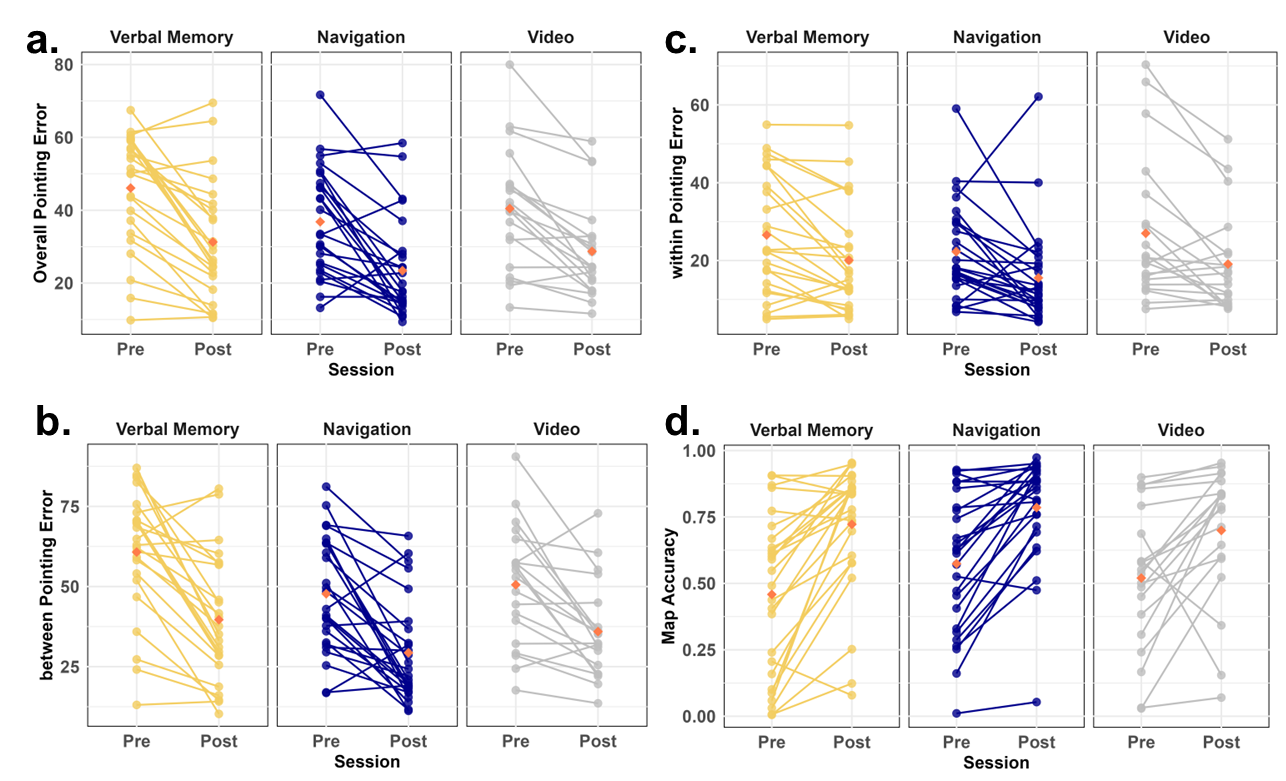

Supplementary Figure 2. Performance on the Navigation Pointing task and Navigation Model Building task.

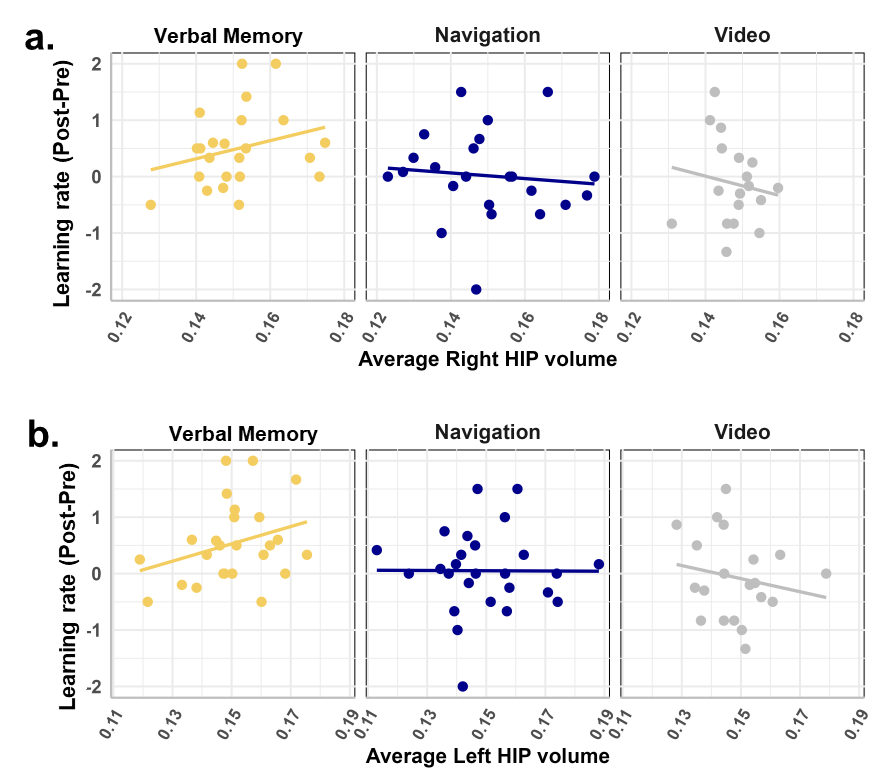

Supplementary Figure 3. Left and right hippocampal volumes were correlated with changes in Verbal Memory Transfer task performance between pre-test and post-test.

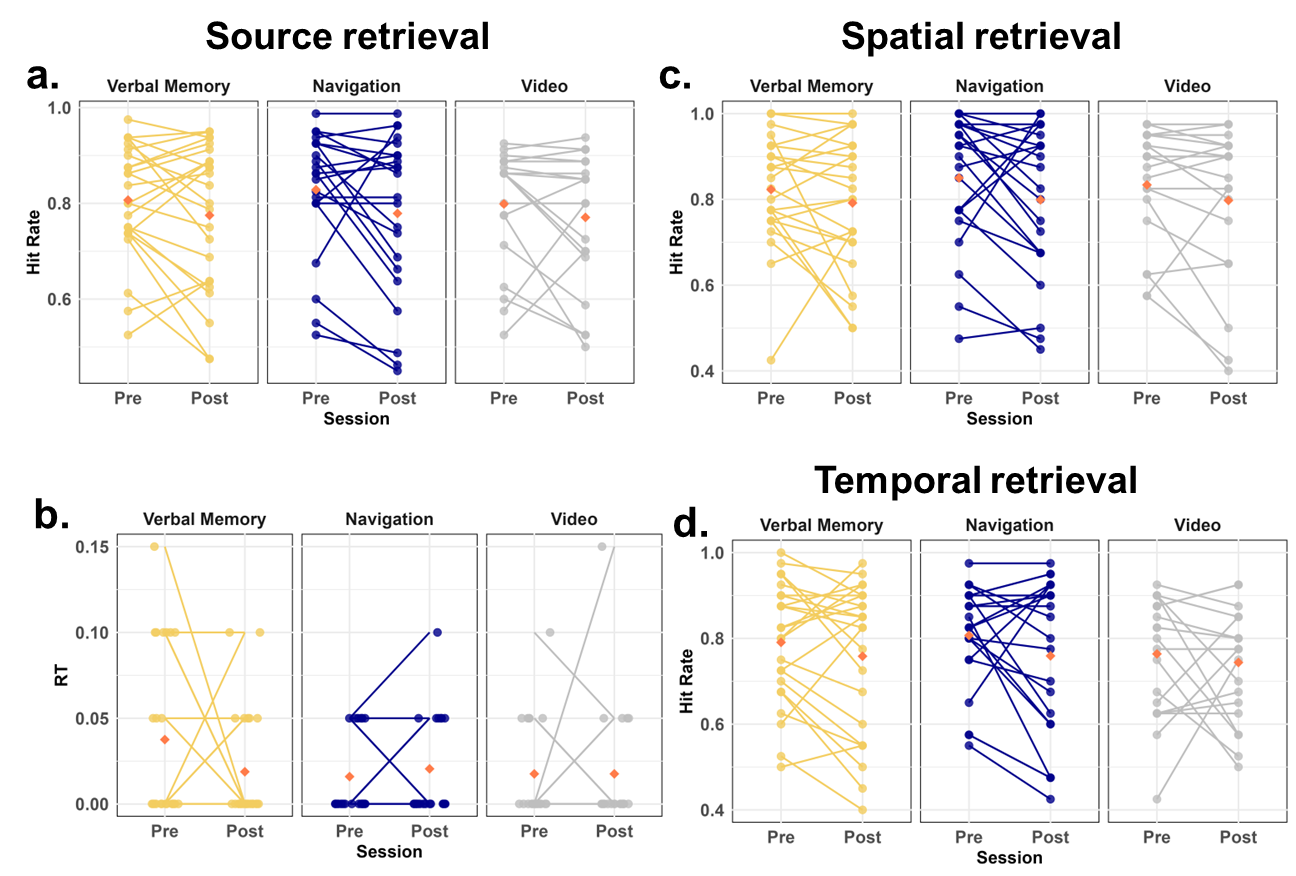

Supplementary Figure 4. Memory performance (hit rate and false alarm rate) during the Source Memory task in the scanner.

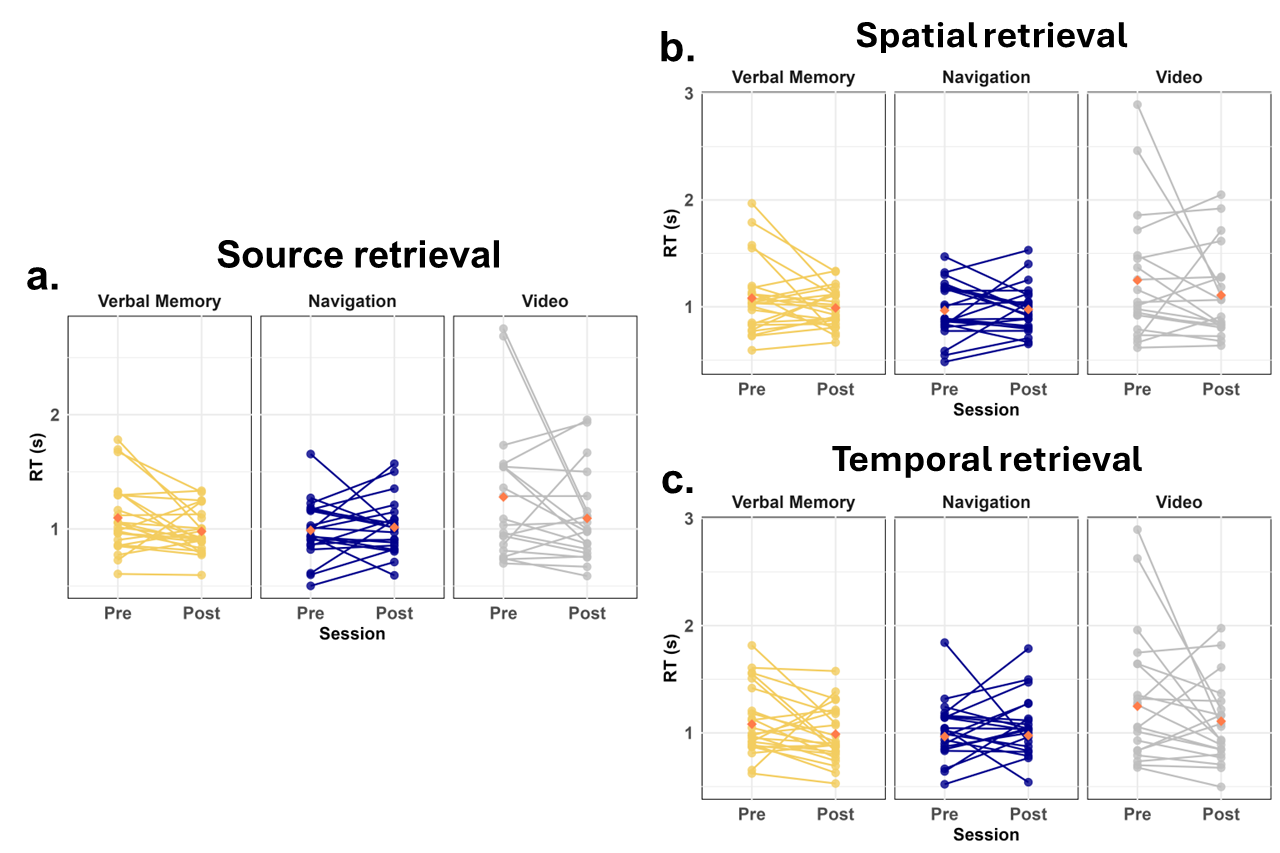

Supplementary Figure 5. Memory performance (reaction time [RT]) during the Source Memory task in the scanner.

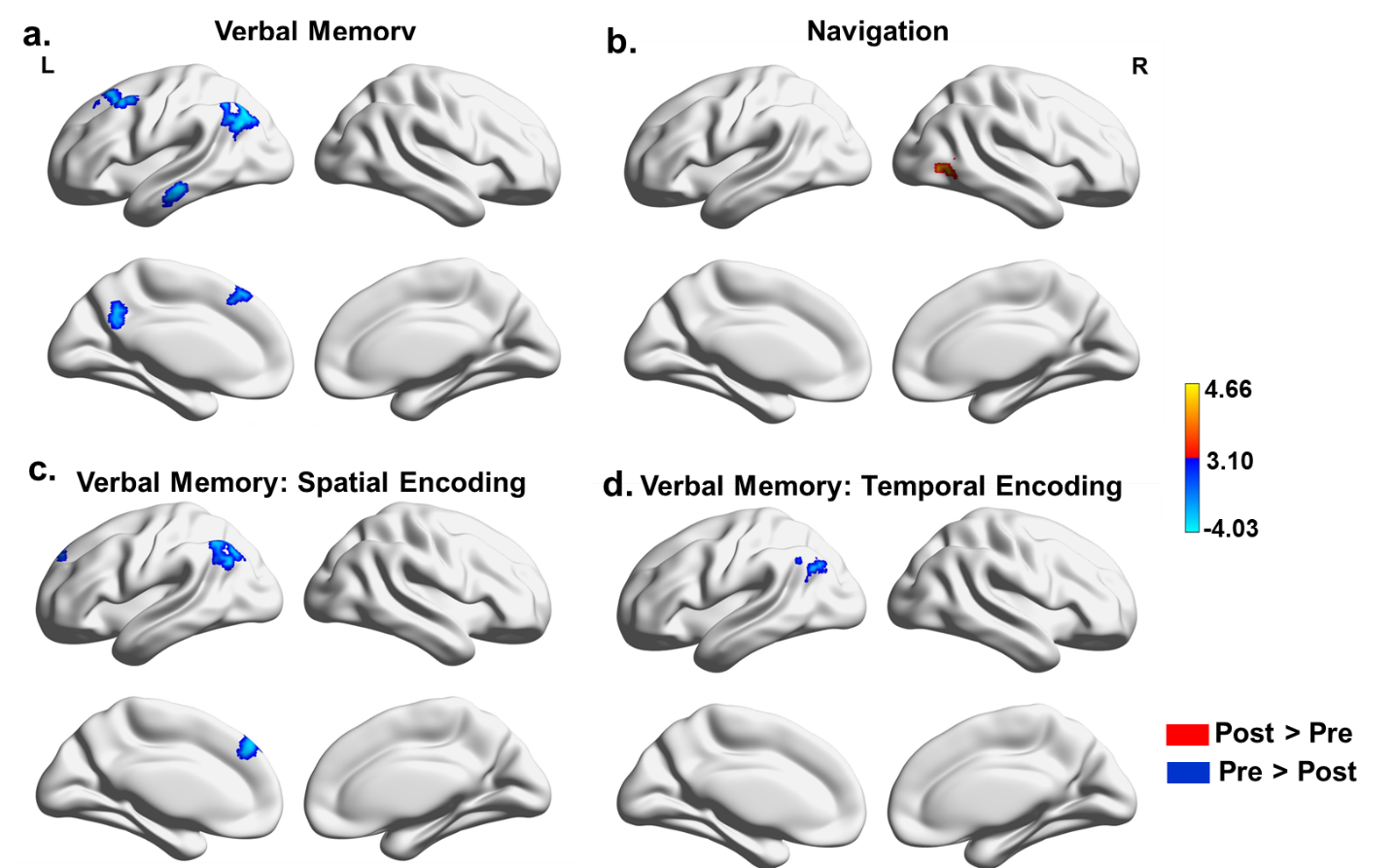

Supplementary Figure 6. Univariate activation changes from pre-test to post-test during Source Memory task encoding.

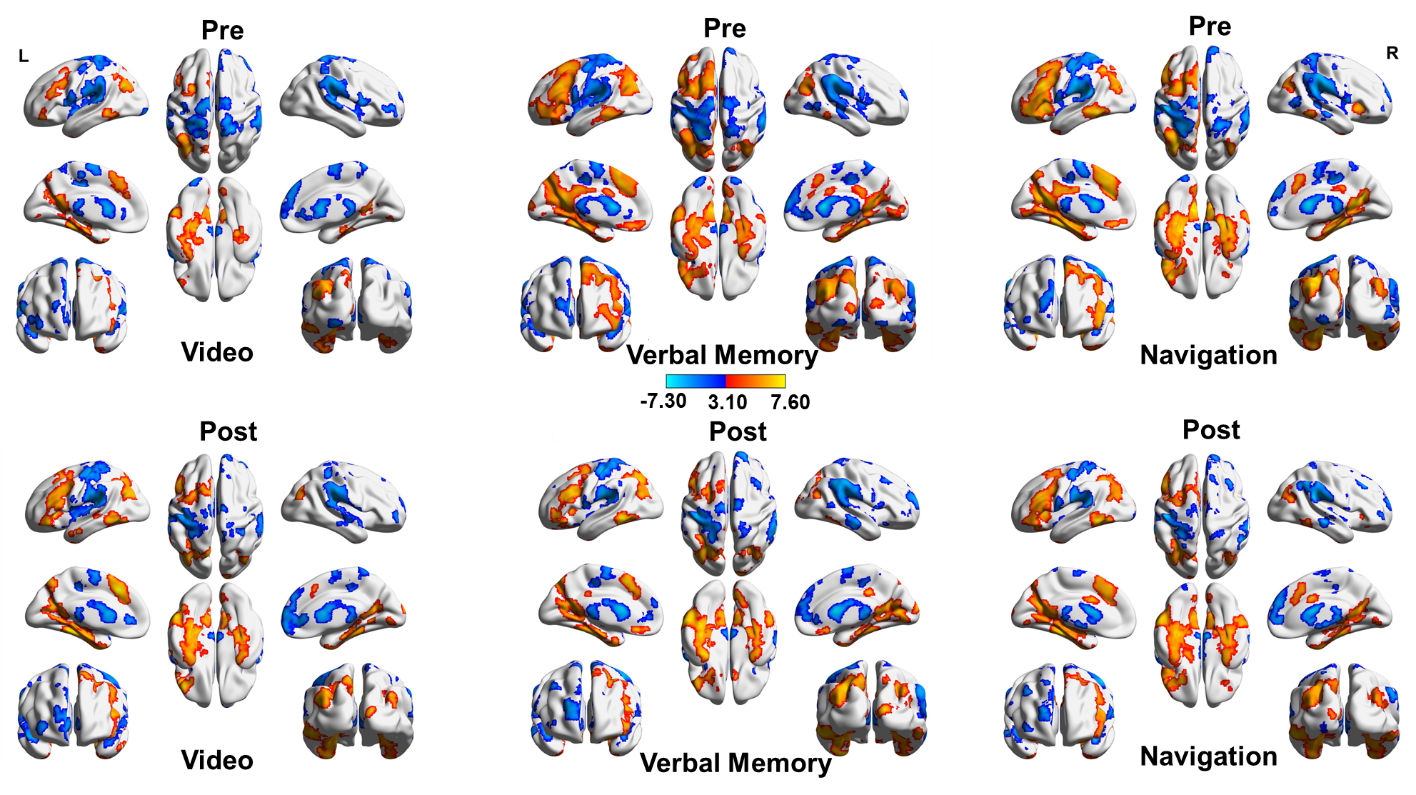

Supplementary Figure 7. Univariate activation during the encoding stage of the Source Memory task, presented separately for pre-test and post-test sessions.

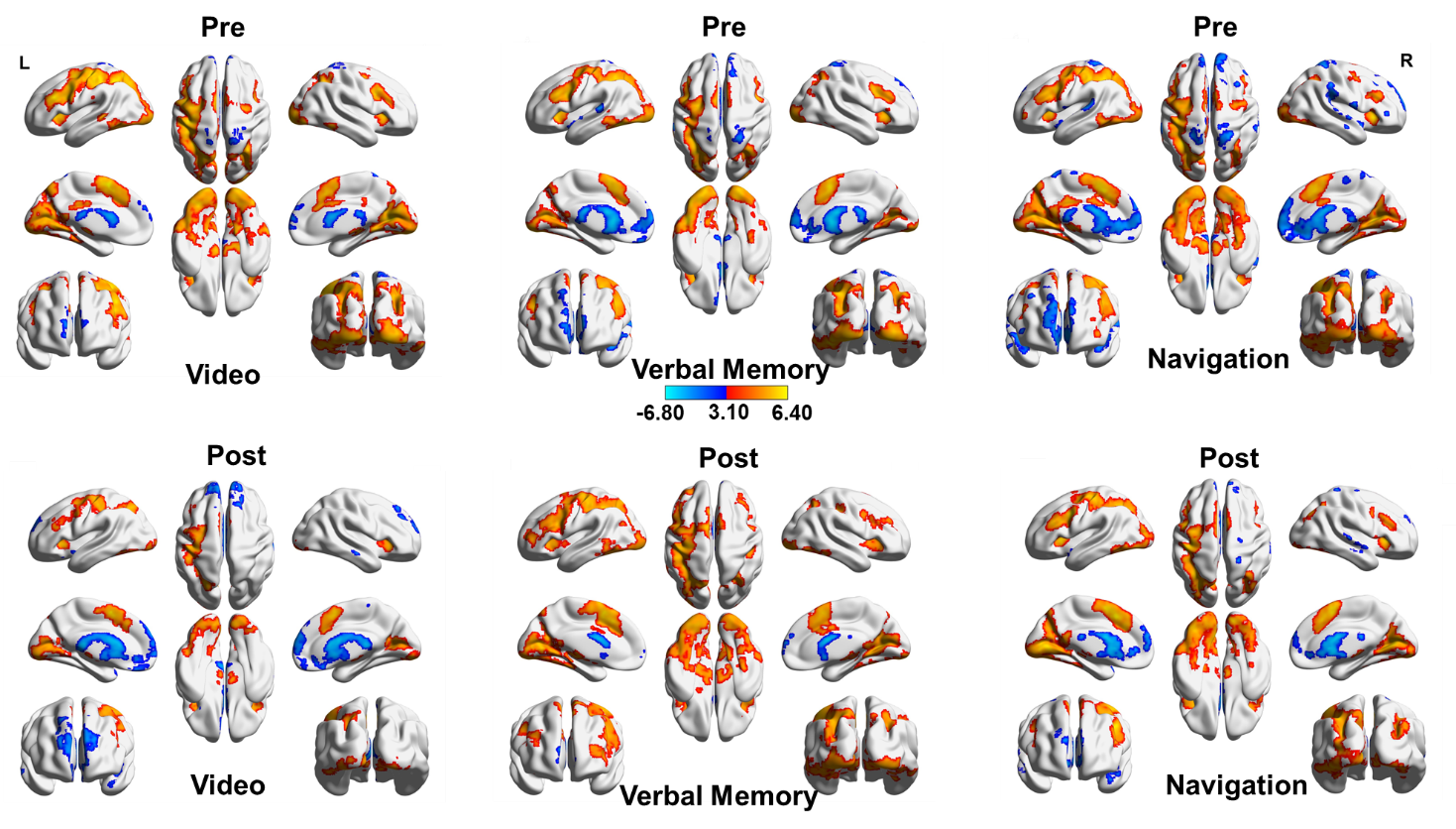

Supplementary Figure 8. Univariate activation during the retrieval stage of the Source Memory task, presented separately for pre-test and post-test sessions.

***Supplementary Tables***

Supplementary Table 1. Demographics information

|  |  | Verbal Memory | Navigation | Video Control | All |
| --- | --- | --- | --- | --- | --- |
|  | Sample size | 27 | 27 | 21 | 75 |
| Sex | Male | 7 | 11 | 7 | 25 |
|  | Female | 20 | 16 | 14 | 50 |
| Site | Site 1 | 13 | 13 | 13 | 39 |
|  | Site 2 | 14 | 14 | 8 | 36 |
|  | Male-Site 1 | 2 | 4 | 5 | 11 |
|  | Female-Site 1 | 9 | 9 | 8 | 26 |
|  | Male-Site 2 | 5 | 7 | 2 | 14 |
|  | Female-Site 2 | 9 | 7 | 6 | 22 |
| Age | Age (min-max/Year) | 20.89 (18-26) | 22.11 (18-32) | 22 (18-32) | 21.67 |
|  | Age-Site1 | 20.31 | 23.00 | 23.00 | 22.10 |
|  | Age-Site2 | 21.43 | 21.29 | 20.38 | 21.03 |

Supplementary Table 2. Sample size information by condition and test, reflecting exclusions due to outlier performance, excessive head movement during scanning, or missing data.

| **Test** | **Verbal Memory** | **Navigation** | **Video** |
| --- | --- | --- | --- |
| Verbal Memory Training | 27 | 27 | 21 |
| Navigation Training | 27 | 27 | 21 |
| Verbal Memory Transfer Task | 26 | 27 | 21 |
| Navigation Transfer Task | 27 | 27 | 19 |
| Navigation Pointing Task | 25 | 27 | 21 |
| Navigation Model Building Task | 27 | 27 | 21 |
| Source Memory Task-Encoding | 25 | 26 | 20 |
| Source Memory Task-Retrieval | 24 | 22 | 20 |
| Hippocampal volume | 26 | 27 | 21 |
| DWI | 20 | 22 | 18 |

Supplementary Table 3. Study Timeline

| **Pre-test**  **(Day 1)** | **Training**  **(Days 2-11)** | **Post-test**  **(Day 12)** |
| --- | --- | --- |
| Complete consent, MRI prescreen, and demographics survey | Comprehensive training varies based on which condition the participant was assigned | Review consent, MRI prescreen |
| Navigation Transfer task (Virtual Silcton, Unity) | Navigation Task (navigation training in virtual Arida, Unity) | Navigation Transfer task (Virtual Silcton, Unity) |
| Navigation Pointing task (Virtual Silcton, web-based) | Verbal memory Task (verbal memory training with the modified method of loci, Unity) | Navigation Pointing task (Virtual Silcton, web-based) |
| Navigation Model Building task (Virtual Silcton, web-based) | Control task (video control by viewing informative videos and answering questions, Qualtrics) | Navigation Model Building task (Virtual Silcton, web-based) |
| Verbal Memory Transfer task (Psychopy) |  | Verbal Memory Transfer task (PsychoPy) |
| Attention task (web-based) |  | Attention task (web-based) |
| Source Memory task (PsychoPy, fMRI scanned) |  | Source Memory task (PsychoPy, fMRI scanned) |
|  |  | Debrief survey (Quatrics) |

Supplementary Table 4. Correlations between the volumes of MTL subregions and changes in learning rate from pre-test to post-test in the Verbal Memory Transfer task.

| Condition | ROI | r | p | Method |
| --- | --- | --- | --- | --- |
| Verbal Memory N=25 | Ant-HIP | 0.248 | 0.232 | Pearson |
|  | Post-HIP | -0.043 | 0.837 | Spearman |
|  | CA1 | 0.138 | 0.51 | Pearson |
|  | CA23DG | 0.532 | **0.006*** | Pearson |
|  | SUB | 0.025 | 0.907 | Pearson |
|  | ERC | 0.069 | 0.743 | Pearson |
|  | PRC | 0.162 | 0.438 | Pearson |
|  | PHC | -0.068 | 0.748 | Spearman |
| Navigation N=27 | Ant-HIP | -0.064 | 0.751 | Spearman |
|  | Post-HIP | 0.183 | 0.362 | Pearson |
|  | CA1 | 0.084 | 0.676 | Pearson |
|  | CA23DG | -0.012 | 0.953 | Pearson |
|  | SUB | 0.035 | 0.863 | Pearson |
|  | ERC | -0.142 | 0.48 | Pearson |
|  | PRC | 0.023 | 0.91 | Pearson |
|  | PHC | 0.223 | 0.264 | Pearson |
| Video N=21 | Ant-HIP | -0.105 | 0.651 | Pearson |
|  | Post-HIP | 0.035 | 0.88 | Pearson |
|  | CA1 | -0.131 | 0.572 | Pearson |
|  | CA23DG | -0.083 | 0.719 | Pearson |
|  | SUB | -0.24 | 0.294 | Pearson |
|  | ERC | 0.149 | 0.52 | Spearman |
|  | PRC | 0.121 | 0.6 | Pearson |
|  | PHC | -0.292 | 0.198 | Pearson |

*Significant results after FDR correction. Ant: anterior; Post: posterior

Supplementary Table 5. Correlations between the volumes of MTL subregions and changes in learning rate from pre-test to post-test in the Navigation Transfer task.

| Condition | ROI | r | p | Method |
| --- | --- | --- | --- | --- |
| Verbal Memory N=26 | Ant-HIP | -0.328 | 0.102 | Pearson |
|  | Post-HIP | -0.147 | 0.471 | Spearman |
|  | HIP | -0.314 | 0.119 | Pearson |
|  | LHIP | -0.291 | 0.149 | Pearson |
|  | RHIP | -0.305 | 0.129 | Pearson |
|  | CA1 | -0.181 | 0.377 | Pearson |
|  | CA23DG | -0.261 | 0.198 | Pearson |
|  | SUB | -0.257 | 0.205 | Pearson |
|  | ERC | 0.027 | 0.897 | Pearson |
|  | PRC | -0.018 | 0.931 | Pearson |
|  | PHC | -0.243 | 0.23 | Spearman |
| Navigation N=27 | Ant-HIP | -0.482 | 0.012 | Spearman |
|  | Post-HIP | -0.042 | 0.837 | Pearson |
|  | HIP | -0.266 | 0.181 | Pearson |
|  | LHIP | -0.247 | 0.214 | Pearson |
|  | RHIP | -0.274 | 0.167 | Pearson |
|  | CA1 | -0.246 | 0.215 | Pearson |
|  | CA23DG | -0.334 | 0.088 | Pearson |
|  | SUB | 0.087 | 0.665 | Pearson |
|  | ERC | 0.008 | 0.967 | Pearson |
|  | PRC | -0.021 | 0.917 | Pearson |
|  | PHC | 0.028 | 0.891 | Pearson |
| Video N=19 | Ant-HIP | -0.101 | 0.681 | Pearson |
|  | Post-HIP | 0.048 | 0.846 | Pearson |
|  | HIP | 0.124 | 0.612 | Pearson |
|  | LHIP | 0.246 | 0.31 | Pearson |
|  | RHIP | 0.028 | 0.911 | Spearman |
|  | CA1 | 0.02 | 0.934 | Pearson |
|  | CA23DG | 0.128 | 0.6 | Pearson |
|  | SUB | 0.053 | 0.828 | Pearson |
|  | ERC | 0.194 | 0.425 | Pearson |
|  | PRC | 0.24 | 0.322 | Pearson |
|  | PHC | -0.089 | 0.716 | Pearson |

Ant: anterior; Post: posterior

Supplementary Table 6. Correlations between the volumes of MTL subregions and changes in the average number of correctly recalled words from pre-test to post-test in the Verbal Memory Transfer task.

| Condition | ROI | r | p | Method |
| --- | --- | --- | --- | --- |
| Verbal Memory N=25 | Ant-HIP | -0.350 | 0.087 | Pearson |
|  | Post-HIP | -0.042 | 0.842 | Spearman |
|  | HIP | -0.235 | 0.258 | Pearson |
|  | LHIP | -0.172 | 0.410 | Pearson |
|  | RHIP | -0.267 | 0.197 | Pearson |
|  | CA1 | -0.212 | 0.309 | Pearson |
|  | CA23DG | -0.329 | 0.109 | Pearson |
|  | SUB | 0.106 | 0.615 | Pearson |
|  | ERC | 0.030 | 0.888 | Pearson |
|  | PRC | -0.219 | 0.292 | Pearson |
|  | PHC | -0.202 | 0.334 | Spearman |
| Navigation N=27 | Ant-HIP | 0.122 | 0.545 | Spearman |
|  | Post-HIP | -0.131 | 0.516 | Pearson |
|  | HIP | 0.012 | 0.953 | Pearson |
|  | LHIP | -0.032 | 0.875 | Pearson |
|  | RHIP | 0.049 | 0.809 | Pearson |
|  | CA1 | 0.023 | 0.911 | Pearson |
|  | CA23DG | 0.043 | 0.830 | Pearson |
|  | SUB | -0.085 | 0.674 | Pearson |
|  | ERC | 0.135 | 0.503 | Pearson |
|  | PRC | -0.062 | 0.757 | Pearson |
|  | PHC | -0.256 | 0.197 | Pearson |
| Video N=21 | Ant-HIP | -0.111 | 0.633 | Pearson |
|  | Post-HIP | -0.012 | 0.959 | Pearson |
|  | HIP | -0.125 | 0.589 | Pearson |
|  | LHIP | -0.156 | 0.499 | Pearson |
|  | RHIP | -0.183 | 0.426 | Spearman |
|  | CA1 | 0.116 | 0.616 | Pearson |
|  | CA23DG | -0.147 | 0.525 | Pearson |
|  | SUB | -0.212 | 0.357 | Pearson |
|  | ERC | -0.365 | 0.104 | Spearman |
|  | PRC | -0.189 | 0.413 | Pearson |
|  | PHC | -0.088 | 0.703 | Pearson |

Ant: anterior; Post: posterior

Supplementary Table 7. Correlations between the volumes of MTL subregions and changes in the number of trials to criterion from pre-test to post-test in the Verbal Memory Transfer task.

| Condition | ROI | r | p | Method |
| --- | --- | --- | --- | --- |
| Verbal Memory N=25 | Ant-HIP | 0.095 | 0.653 | Spearman |
|  | Post-HIP | 0.316 | 0.124 | Spearman |
|  | HIP | -0.163 | 0.438 | Spearman |
|  | LHIP | -0.111 | 0.598 | Spearman |
|  | RHIP | -0.223 | 0.285 | Spearman |
|  | CA1 | 0.028 | 0.896 | Spearman |
|  | CA23DG | 0.074 | 0.726 | Spearman |
|  | SUB | -0.398 | 0.049 | Spearman |
|  | ERC | -0.334 | 0.103 | Spearman |
|  | PRC | 0.139 | 0.509 | Spearman |
|  | PHC | 0.041 | 0.846 | Spearman |
| Navigation N=27 | Ant-HIP | 0.084 | 0.679 | Spearman |
|  | Post-HIP | 0.008 | 0.970 | Spearman |
|  | HIP | -0.023 | 0.908 | Spearman |
|  | LHIP | -0.042 | 0.837 | Spearman |
|  | RHIP | -0.033 | 0.872 | Spearman |
|  | CA1 | -0.015 | 0.943 | Spearman |
|  | CA23DG | -0.028 | 0.890 | Spearman |
|  | SUB | 0.099 | 0.622 | Spearman |
|  | ERC | -0.011 | 0.956 | Spearman |
|  | PRC | 0.137 | 0.495 | Spearman |
|  | PHC | 0.257 | 0.196 | Spearman |
| Video N=21 | Ant-HIP | 0.223 | 0.332 | Pearson |
|  | Post-HIP | 0.242 | 0.290 | Pearson |
|  | HIP | 0.359 | 0.110 | Pearson |
|  | LHIP | 0.413 | 0.063 | Pearson |
|  | RHIP | 0.222 | 0.334 | Spearman |
|  | CA1 | 0.167 | 0.471 | Pearson |
|  | CA23DG | 0.178 | 0.440 | Pearson |
|  | SUB | 0.436 | 0.048 | Pearson |
|  | ERC | 0.202 | 0.379 | Spearman |
|  | PRC | 0.072 | 0.756 | Pearson |
|  | PHC | 0.461 | 0.035 | Pearson |

Ant: anterior; Post: posterior

Supplementary Table 8. Correlations between the volumes of MTL subregions and changes in slope from pre-test to post-test in the Verbal Memory Transfer task.

| Condition | ROI | r | p | Method |
| --- | --- | --- | --- | --- |
| Verbal Memory N=26 | Ant-HIP | 0.30 | 0.15 | Pearson |
|  | Post-HIP | -0.14 | 0.51 | Spearman |
|  | HIP | 0.39 | 0.05 | Pearson |
|  | LHIP | 0.32 | 0.12 | Pearson |
|  | RHIP | 0.42 | 0.04 | Pearson |
|  | CA1 | 0.13 | 0.54 | Pearson |
|  | CA23DG | 0.50 | 0.01 | Pearson |
|  | SUB | 0.15 | 0.46 | Pearson |
|  | ERC | 0.11 | 0.59 | Pearson |
|  | PRC | 0.19 | 0.37 | Pearson |
|  | PHC | 0.01 | 0.95 | Spearman |
| Navigation N=27 | Ant-HIP | 0.04 | 0.83 | Spearman |
|  | Post-HIP | 0.23 | 0.26 | Pearson |
|  | HIP | 0.09 | 0.67 | Pearson |
|  | LHIP | 0.10 | 0.63 | Pearson |
|  | RHIP | 0.07 | 0.71 | Pearson |
|  | CA1 | 0.18 | 0.36 | Pearson |
|  | CA23DG | 0.02 | 0.92 | Pearson |
|  | SUB | 0.06 | 0.75 | Pearson |
|  | ERC | -0.10 | 0.63 | Pearson |
|  | PRC | 0.01 | 0.97 | Pearson |
|  | PHC | 0.21 | 0.30 | Pearson |
| Video N=21 | Ant-HIP | -0.23 | 0.32 | Pearson |
|  | Post-HIP | -0.05 | 0.82 | Pearson |
|  | HIP | -0.29 | 0.20 | Pearson |
|  | LHIP | -0.28 | 0.21 | Pearson |
|  | RHIP | -0.28 | 0.21 | Spearman |
|  | CA1 | -0.16 | 0.50 | Pearson |
|  | CA23DG | -0.15 | 0.51 | Pearson |
|  | SUB | -0.31 | 0.17 | Pearson |
|  | ERC | 0.09 | 0.71 | Spearman |
|  | PRC | 0.05 | 0.82 | Pearson |
|  | PHC | -0.33 | 0.14 | Pearson |

Ant: anterior; Post: posterior

Supplementary Table 9. Correlations between the volumes of MTL subregions and changes in path errors from pre-test to post-test in the Navigation Transfer task.

| Condition | ROI | r | p | Method |
| --- | --- | --- | --- | --- |
| Verbal Memory N=26 | Ant-HIP | 0.080 | 0.698 | Pearson |
|  | Post-HIP | 0.117 | 0.570 | Spearman |
|  | HIP | 0.041 | 0.844 | Pearson |
|  | LHIP | 0.117 | 0.570 | Pearson |
|  | RHIP | 0.041 | 0.844 | Pearson |
|  | CA1 | 0.163 | 0.426 | Pearson |
|  | CA23DG | -0.214 | 0.294 | Pearson |
|  | ERC | 0.274 | 0.176 | Pearson |
|  | PHC | 0.047 | 0.820 | Pearson |
|  | PRC | -0.452 | 0.020 | Pearson |
|  | SUB | 0.396 | 0.045 | Spearman |
| Navigation N=27 | Ant-HIP | -0.137 | 0.493 | Spearman |
|  | Post-HIP | -0.181 | 0.366 | Pearson |
|  | HIP | -0.058 | 0.775 | Pearson |
|  | LHIP | -0.032 | 0.873 | Pearson |
|  | RHIP | -0.078 | 0.701 | Pearson |
|  | CA1 | -0.294 | 0.137 | Pearson |
|  | CA23DG | 0.011 | 0.956 | Pearson |
|  | ERC | -0.072 | 0.723 | Pearson |
|  | PHC | 0.027 | 0.892 | Pearson |
|  | PRC | -0.083 | 0.681 | Pearson |
|  | SUB | 0.101 | 0.617 | Pearson |
| Video N=19 | Ant-HIP | 0.301 | 0.211 | Pearson |
|  | Post-HIP | 0.210 | 0.389 | Pearson |
|  | HIP | -0.193 | 0.429 | Pearson |
|  | LHIP | -0.187 | 0.443 | Pearson |
|  | RHIP | -0.130 | 0.595 | Spearman |
|  | CA1 | -0.186 | 0.446 | Pearson |
|  | CA23DG | 0.052 | 0.833 | Pearson |
|  | ERC | 0.065 | 0.791 | Pearson |
|  | PHC | -0.291 | 0.227 | Pearson |
|  | PRC | 0.150 | 0.540 | Pearson |
|  | SUB | -0.277 | 0.252 | Pearson |

Ant: anterior; Post: posterior

Supplementary Table 10. Correlations between the volumes of MTL subregions and changes in overall pointing errors from pre-test to post-test in the Navigation Pointing Error task.

| Condition | ROI | r | p | Method |
| --- | --- | --- | --- | --- |
| Verbal Memory N=24 | Ant-HIP | -0.143 | 0.504 | Pearson |
|  | Post-HIP | 0.191 | 0.369 | Spearman |
|  | HIP | -0.205 | 0.336 | Pearson |
|  | LHIP | -0.127 | 0.554 | Pearson |
|  | RHIP | -0.252 | 0.236 | Pearson |
|  | CA1 | 0.000 | 1.000 | Pearson |
|  | CA23DG | -0.201 | 0.346 | Pearson |
|  | ERC | -0.086 | 0.690 | Pearson |
|  | PHC | -0.134 | 0.531 | Spearman |
|  | PRC | 0.314 | 0.136 | Pearson |
|  | SUB | -0.263 | 0.214 | Pearson |
| Navigation N=27 | Ant-HIP | -0.119 | 0.553 | Spearman |
|  | Post-HIP | -0.317 | 0.107 | Pearson |
|  | HIP | -0.159 | 0.429 | Pearson |
|  | LHIP | -0.198 | 0.322 | Pearson |
|  | RHIP | -0.121 | 0.548 | Pearson |
|  | CA1 | -0.295 | 0.136 | Pearson |
|  | CA23DG | -0.017 | 0.932 | Pearson |
|  | ERC | -0.024 | 0.907 | Pearson |
|  | PHC | -0.227 | 0.256 | Pearson |
|  | PRC | -0.090 | 0.656 | Pearson |
|  | SUB | -0.208 | 0.299 | Pearson |
| Video N=21 | Ant-HIP | -0.155 | 0.503 | Pearson |
|  | Post-HIP | 0.041 | 0.860 | Pearson |
|  | HIP | 0.200 | 0.385 | Pearson |
|  | LHIP | 0.247 | 0.280 | Pearson |
|  | RHIP | 0.042 | 0.859 | Spearman |
|  | CA1 | -0.040 | 0.864 | Pearson |
|  | CA23DG | 0.188 | 0.415 | Pearson |
|  | ERC | -0.246 | 0.282 | Spearman |
|  | PHC | 0.247 | 0.282 | Pearson |
|  | PRC | 0.238 | 0.300 | Pearson |
|  | SUB | 0.250 | 0.274 | Pearson |

Ant: anterior; Post: posterior

Supplementary Table 11. Correlations between the volumes of MTL subregions and changes in within-environment pointing error from pre-test to post-test in the Navigation Pointing Error task.

| Condition | ROI | r | | p | Method |
| --- | --- | --- | --- | --- | --- |
| Verbal Memory N=24 | Ant-HIP | -0.142 | | 0.508 | Pearson |
|  | Post-HIP | | 0.165 | 0.439 | Spearman |
|  | HIP | 0.110 | | 0.610 | Pearson |
|  | LHIP | 0.163 | | 0.447 | Pearson |
|  | RHIP | 0.055 | | 0.798 | Pearson |
|  | CA1 | 0.040 | | 0.854 | Pearson |
|  | CA23DG | 0.239 | | 0.261 | Pearson |
|  | ERC | 0.192 | | 0.369 | Pearson |
|  | PHC | -0.145 | | 0.497 | Spearman |
|  | PRC | 0.411 | | 0.046 | Pearson |
|  | SUB | -0.149 | | 0.488 | Pearson |
| Navigation N=27 | Ant-HIP | -0.242 | | 0.222 | Spearman |
|  | Post-HIP | -0.249 | | 0.211 | Pearson |
|  | HIP | -0.166 | | 0.408 | Pearson |
|  | LHIP | -0.183 | | 0.360 | Pearson |
|  | RHIP | -0.147 | | 0.466 | Pearson |
|  | CA1 | -0.199 | | 0.321 | Pearson |
|  | CA23DG | -0.160 | | 0.427 | Pearson |
|  | ERC | 0.019 | | 0.925 | Pearson |
|  | PHC | -0.178 | | 0.374 | Pearson |
|  | PRC | -0.204 | | 0.309 | Pearson |
|  | SUB | -0.010 | | 0.961 | Pearson |
| Video N=21 | Ant-HIP | 0.166 | | 0.471 | Pearson |
|  | Post-HIP | 0.181 | | 0.432 | Pearson |
|  | HIP | 0.060 | | 0.796 | Pearson |
|  | LHIP | 0.040 | | 0.865 | Pearson |
|  | RHIP | 0.044 | | 0.850 | Spearman |
|  | CA1 | -0.078 | | 0.738 | Pearson |
|  | CA23DG | 0.301 | | 0.185 | Pearson |
|  | ERC | -0.260 | | 0.254 | Spearman |
|  | PHC | -0.112 | | 0.629 | Pearson |
|  | PRC | 0.415 | | 0.062 | Pearson |
|  | SUB | -0.232 | | 0.312 | Pearson |

Ant: anterior; Post: posterior

Supplementary Table 12. Correlations between the volumes of MTL subregions and changes in between-environment pointing error from pre-test to post-test in the Navigation Pointing Error Task.

| Condition | ROI | r | p | Method |
| --- | --- | --- | --- | --- |
| Verbal Memory N=24 | Ant-HIP | -0.120 | 0.576 | Pearson |
|  | Post-HIP | 0.190 | 0.373 | Spearman |
|  | HIP | -0.285 | 0.177 | Pearson |
|  | LHIP | -0.211 | 0.323 | Pearson |
|  | RHIP | -0.321 | 0.127 | Pearson |
|  | CA1 | -0.014 | 0.948 | Pearson |
|  | CA23DG | -0.327 | 0.119 | Pearson |
|  | ERC | -0.172 | 0.423 | Pearson |
|  | PHC | -0.183 | 0.391 | Spearman |
|  | PRC | 0.227 | 0.286 | Pearson |
|  | SUB | -0.261 | 0.218 | Pearson |
| Navigation N=27 | Ant-HIP | -0.145 | 0.468 | Spearman |
|  | Post-HIP | -0.295 | 0.136 | Pearson |
|  | HIP | -0.125 | 0.534 | Pearson |
|  | LHIP | -0.169 | 0.400 | Pearson |
|  | RHIP | -0.084 | 0.676 | Pearson |
|  | CA1 | -0.291 | 0.141 | Pearson |
|  | CA23DG | 0.063 | 0.756 | Pearson |
|  | ERC | -0.042 | 0.834 | Pearson |
|  | PHC | -0.210 | 0.294 | Pearson |
|  | PRC | -0.011 | 0.955 | Pearson |
|  | SUB | -0.275 | 0.165 | Pearson |
| Video N=21 | Ant-HIP | -0.248 | 0.278 | Pearson |
|  | Post-HIP | -0.047 | 0.838 | Pearson |
|  | HIP | 0.182 | 0.430 | Pearson |
|  | LHIP | 0.243 | 0.290 | Pearson |
|  | RHIP | -0.021 | 0.930 | Spearman |
|  | CA1 | -0.003 | 0.989 | Pearson |
|  | CA23DG | 0.049 | 0.835 | Pearson |
|  | ERC | -0.183 | 0.425 | Spearman |
|  | PHC | 0.318 | 0.160 | Pearson |
|  | PRC | 0.044 | 0.850 | Pearson |
|  | SUB | 0.382 | 0.087 | Pearson |

Ant: anterior; Post: posterior

Supplementary Table 13. Correlations between the volumes of MTL subregions and changes in model-building accuracy from pre-test to post-test in the Navigation Model Building task.

| Condition | ROI | r | p | Method |
| --- | --- | --- | --- | --- |
| Verbal Memory N=26 | Ant-HIP | 0.125 | 0.544 | Pearson |
|  | Post-HIP | -0.158 | 0.438 | Spearman |
|  | HIP | 0.066 | 0.749 | Pearson |
|  | LHIP | 0.025 | 0.905 | Pearson |
|  | RHIP | 0.096 | 0.640 | Pearson |
|  | CA1 | -0.048 | 0.816 | Pearson |
|  | CA23DG | 0.137 | 0.504 | Pearson |
|  | ERC | -0.009 | 0.965 | Pearson |
|  | PHC | 0.074 | 0.721 | Spearman |
|  | PRC | 0.345 | 0.085 | Pearson |
|  | SUB | 0.017 | 0.936 | Pearson |
| Navigation N=27 | Ant-HIP | -0.035 | 0.863 | Spearman |
|  | Post-HIP | 0.008 | 0.969 | Pearson |
|  | HIP | 0.078 | 0.701 | Pearson |
|  | LHIP | 0.046 | 0.821 | Pearson |
|  | RHIP | 0.102 | 0.612 | Pearson |
|  | CA1 | 0.322 | 0.102 | Pearson |
|  | CA23DG | -0.068 | 0.737 | Pearson |
|  | ERC | -0.065 | 0.746 | Pearson |
|  | PHC | -0.038 | 0.852 | Pearson |
|  | PRC | -0.129 | 0.520 | Pearson |
|  | SUB | 0.074 | 0.715 | Pearson |
| Video N=21 | Ant-HIP | -0.083 | 0.722 | Pearson |
|  | Post-HIP | -0.231 | 0.313 | Pearson |
|  | HIP | 0.039 | 0.868 | Pearson |
|  | LHIP | -0.020 | 0.931 | Pearson |
|  | RHIP | 0.092 | 0.690 | Spearman |
|  | CA1 | 0.129 | 0.577 | Pearson |
|  | CA23DG | -0.181 | 0.432 | Pearson |
|  | ERC | 0.203 | 0.377 | Spearman |
|  | PHC | 0.046 | 0.842 | Pearson |
|  | PRC | -0.407 | 0.067 | Pearson |
|  | SUB | 0.235 | 0.304 | Pearson |

Ant: anterior; Post: posterior

Supplementary Table 14. Correlations between the average volumes of MTL subregions and slope from Day 1 to Day 5 in the Verbal Memory Training task.

| Condition | ROI | r | p | Method |
| --- | --- | --- | --- | --- |
| Verbal Memory N=26 | Ant-HIP | -0.0323 | 0.8755 | Pearson |
|  | Post-HIP | -0.1321 | 0.52 | Spearman |
|  | HIP | 0.0912 | 0.6577 | Pearson |
|  | LHIP | 0.2391 | 0.2395 | Pearson |
|  | RHIP | -0.0455 | 0.8252 | Pearson |
|  | CA1 | -0.0993 | 0.6292 | Pearson |
|  | CA23DG | 0.0299 | 0.8846 | Pearson |
|  | ERC | 0.1806 | 0.3773 | Pearson |
|  | PHC | 0.1397 | 0.4962 | Spearman |
|  | PRC | -0.3257 | 0.1044 | Pearson |
|  | SUB | 0.3239 | 0.1065 | Pearson |

Ant: anterior; Post: posterior

Supplementary Table 15. Correlations between the average volumes of MTL subregions and slope from Day 6 to Day 10 in the Verbal Memory Training task.

| Condition | ROI | r | p | Method |
| --- | --- | --- | --- | --- |
| Verbal Memory N=26 | Ant-HIP | -0.0882 | 0.6682 | Pearson |
|  | Post-HIP | -0.0576 | 0.78 | Spearman |
|  | HIP | -0.1188 | 0.5633 | Pearson |
|  | LHIP | 0.0082 | 0.9685 | Pearson |
|  | RHIP | -0.2188 | 0.2829 | Pearson |
|  | CA1 | -0.0902 | 0.6612 | Pearson |
|  | CA23DG | -0.2916 | 0.1484 | Pearson |
|  | ERC | 0.1524 | 0.4573 | Pearson |
|  | PHC | 0.1542 | 0.4521 | Spearman |
|  | PRC | -0.3393 | 0.09 | Pearson |
|  | SUB | 0.2459 | 0.2259 | Pearson |

Ant: anterior; Post: posterior

Supplementary Table 16. Univariate activation changes from pre-test to post-test during encoding and retrieval, with whole-brain cluster correction and small-volume correction for the hippocampus reported separately.

| **Pre vs. Post** | | | |
| --- | --- | --- | --- |
| **Condition** | **Contrast** | **Results (Hippocampus)** | **Results**  **(whole brain)** |
| Encoding | Video | None | None |
|  | Verbal Memory | None | T1 > T2:  left dLOC (Z = 4.66, MNI: -46, -62, 30), left MFG (Z = 4.32, MNI: -42, 12, 50), left frontal pole (Z = 4.2, MNI: -24, 40, 46), left MTG (Z = 4.28, MNI: -60, -24, -12), left precuneus (Z = 3.8, MNI: -10, -52, 38). |
|  | Navigation | None | T2 > T1:  right MTG (Z = 4.03, MNI: 42, -54, ss6). |
|  | Verbal Memory vs. Video | None | None |
|  | Navigation vs. Video | None | None |
|  | Verbal Memory vs. Navigation | None | None |
|  | Verbal Memory vs. Video+Navigation | None | None |
|  | Navigation vs. Video+Verbal Memory | None | None |
| Spatial encoding | Video | None | None |
|  | Verbal Memory | None | T1 > T2: Left LOC (Z = 3.97, MNI: -44,-62,32), Left frontal pole (Z = 3.99, MNI: -18, 48, 36). |
|  | Navigation | None | None |
|  | Verbal Memory vs. Video | None | None |
|  | Navigation vs. Video | None | None |
|  | Verbal Memory vs. Navigation | None | None |
|  | Verbal Memory vs. Video + Navigation | None | None |
|  | Navigation vs. Video + Verbal Memory | None | None |
| Temporal encoding | Video | None | None |
|  | Verbal Memory | None | T1 > T2: Left LOC (Z = 3.97, MNI: -40, -62, 32) |
|  | Navigation | None | None |
|  | Verbal Memory vs. Video | None | None |
|  | Navigation vs. Video | None | None |
|  | Verbal Memory vs. Navigation | None | None |
|  | Verbal Memory vs. Video + Navigation | None | None |
|  | Navigation vs. Video + Verbal Memory | None | None |
| Retrieval | Video | None | None |
|  | Verbal Memory | None | None |
|  | Navigation | None | None |
|  | Verbal Memory vs. Video | None | None |
|  | Navigation vs. Video | None | None |
|  | Verbal Memory vs. Navigation | None | None |
|  | Verbal Memory vs. Video + Navigation | None | None |
|  | Navigation vs. Video + Verbal Memory | None | None |
| Spatial Retrieval | Video | None | None |
|  | Verbal Memory | None | None |
|  | Navigation | None | None |
|  | Verbal Memory vs. Video | None | None |
|  | Navigation vs. Video | None | None |
|  | Verbal Memory vs. Navigation | None | None |
|  | Verbal Memory vs. Video + Navigation | None | None |
|  | Navigation vs. Video + Verbal Memory | None | None |
| Temporal Retrieval | Video | None | None |
|  | Verbal Memory | None | None |
|  | Navigation | None | None |
|  | Verbal Memory vs. Video | None | None |
|  | Navigation vs. Video | None | None |
|  | Verbal Memory vs. Navigation | None | None |
|  | Verbal Memory vs. Video + Navigation | None | None |
|  | Navigation vs. Video + Verbal Memory | None | None |

**Supplementary Notes**

**Supplementary Notes 1: Methods**

***Demographic Survey***

Participants completed an online series of self-report surveys including demographic information (i.e., age, sex at birth, gender, ethnicity, race), MRI survey (handedness, color vision, visual acuity), virtual reality/cybersickness survey (number of hours they slept the night prior, last meal, virtual reality exposure, video game experience, weight, history of motion sickness, history of dizziness/fainting/syncope), and other pertinent study information (level of stress, source of stress).

***Pre-post tests***

***Navigation Pointing Task: Virtual Silcton***

Using the Virtual Silcton website (<http://www.virtualsilcton.com/>), participants were placed at the goal location in front of each of the 8 buildings within the Virtual Silcton environment. Participants were not capable of translation in this portion of the task, but fully capable of rotation. Participants were told to point the target on the screen in the direction of the other 7 target buildings, then click the mouse to record their response (see Supplementary Figure 1a).

***Navigation Model Building Task: Virtual Silcton***

Continuing on the Virtual Silcton website, participants were presented with a blank box and instructed that this box represents the boundaries of the Virtual Silcton environment. To the right of the square, they were shown bird’s-eye view images of the 8 destinations from the Navigation Task. Using the mouse, they dragged and dropped each building into the blank square to create a map of how they recalled the virtual environment (see Supplementary Figure 1b). Once finished, participants clicked a button that indicated that they had completed the task.

***Attention Task***

Participants completed a Posner cueing task (Posner, 1980) in which they were told to respond to an appearance of a star that would either appear to the left or to the right of a fixation cross through valid and invalid highlighting cues. Participants were instructed to be accurate, but quick. This task was completed via the Testable website (<https://www.testable.org/project>).

**Supplementary Note 2: Training-related effects**

*Control analysis for the Navigation task* (*navigation training in virtual Arida) .* Regarding average distance error traveled for individual subsections, we observed a significant main effect of subsection *F*(2,42) = 13.46, *p* < .001, η^2^ = .39. We also observed significant effects of sex and site (men outperformed women and participants at the University of Arizona outperformed participants at the University of Florida), but these did not interact with each other, nor with the within-participant training effects (*p*s > .41). Removing sex and site from the model did not change the effect. Removing outliers (3 standard deviations away from the grand mean) did not affect results. Regarding integrating multiple subsections, we observed a significant main effect of subsections, *F*(1,22) = 5.16, *p* = .03, η^2^ = .19. This finding was more fragile. Including sex and site in the model reduced the finding to non-significant (*p* = .06). This analysis was also not robust to outlier removal.

*Control analysis for Verbal Memory task (verbal memory training) performance.* Significant improvements in verbal memory performance were observed throughout 10 days training, with slopes significantly greater than zero after controlling for sex and site for the first five days (t(26) = 3.964, p = 0.001) and the last five days (t(26) = 4.877, p < .001).

**Supplementary Note 3: Pre-test and post-test behavioral transfer**

***Verbal Memory Transfer task***

*Control analysis for Learning rate of Verbal Memory Transfer task*

When controlling for sex and site using linear mixed effect model, the Verbal Memory group still showed an increase in learning rate from pre-test to post-test (t(26) = 3.381, p = 0.002, Marginal R^2^ = 0.117) compared to the Navigation group (t(27) = 0.497, p = 0.623, Marginal R^2^ = 0.016) and Video Control group (t(21) = 0.564, p = 0.588, Marginal R^2^ = 0.025).

In further analysis of the learning rate for the Verbal Memory Transfer task, a potential issue arises with participants who met the criteria within a single trial, making it problematic to calculate the difference between the last and first trials. In the initial analysis, these participants were assigned a value of zero. However, in the subsequent control analysis, we excluded these participants, one from the Video Control group and two from the Navigation group, all of whom met the criteria within one trial in the post-test. All results remained consistent after excluding those participants: A mixed-design ANOVA, with condition (Navigation/Verbal Memory/Video Control) as a between-subjects factor and session (pre/post) as a within-subjects factor, revealed a main effect of session (F(1, 68) = 6.724, p = 0.012, η^2^ = .028) and a significant interaction between condition and session (F(2, 68) = 3.410, p = 0.039, η^2^ = .028). Paired-sample t-tests indicated that only the Verbal Memory group showed a significant increase in learning rate from pre to post (Verbal Memory: t(25) = 3.316, p = 0.003, Cohen’s d = 0.608; Navigation: t(24) = 1.697, p = 0.103, Cohen’s d = 0.347; Video Control: t(19) = -0.263, p = 0.795, Cohen’s d = -0.081). When controlling for sex and site in linear mixed effect models, the Verbal Memory group still showed the greatest increase in learning rate from pre to post (t(26) = 3.381, p = 0.002, Marginal R^2^ = 0.117) compared to the Navigation group (t(25) = 1.732, p = 0.096, Marginal R^2^ = 0.06) and Video Control group (t(20) = 0.270, p = 0.790, Marginal R^2^ = 0.007).

*Slope of Verbal Memory Transfer task*

In a complementary analysis, we calculated the slope across trials as an alternative measure of the learning rate and observed similar results. The slope was derived using linear regression on the number of words recalled in each trial. A mixed-design ANOVA, with condition (Navigation/Verbal Memory/Video Control) as a between-subjects factor and session (pre/post) as a within-subjects factor, revealed a main effect of session (F(1, 71) = 7.144, p = 0.009, η^2^ = .032) but no significant interaction between condition and session (F(2, 71) = 2.579, p = 0.083, η^2^ = .023). Paired-sample t-tests indicated that only the Verbal Memory group showed a significant increase in learning rate from pre to post (Verbal Memory: t(25) = 3.56, p = 0.002, Cohen’s d = 0.631; Navigation: t(26) = 0.909, p = 0.372, Cohen’s d = 0.212; Video Control: t(20) = 0.369, p = 0.716, Cohen’s d = -0.121). When controlling for sex and site in linear mixed effect models, the Verbal Memory group still showed a significant increase in learning rate from pre-test to post-test (t(26) = 3.627, p = 0.004, Marginal R^2^ = 0.135) compared to the Navigation group (t(27) = 0.926, p = 0.362, Marginal R^2^ = 0.075) and Video Control group (t(38) = 0.384, p = 0.703, partial η^2^ = 0.0039).

*During the pre-test session and post-test session, we also calculated classic indices of verbal memory performance, including the average number of words recalled for the 12-word list and the number of trials required to reach the criterion (i.e., 12) in the Verbal Memory Transfer task.*

A mixed-design ANOVA was conducted with average number of words correctly recalled as the dependent variable, condition (Navigation/Verbal Memory/Video Control) as a between-subjects factor, and session (pre/post) as a within-subjects factor. The analysis revealed a significant interaction between condition and session (F(2, 71) = 8.163, p < 0.001, η^2^ = .0.049). Surprisingly, paired-sample t-tests showed that only the Video Control group demonstrated a significant increase in average number of words recalled from pre-test to post-test (t(20) = 3.28, p = 0.004, Cohen’s d = 0.69). In contrast, the Verbal Memory group exhibited a significant decrease in average number of words recalled (t(25) = -2.26, p = 0.033, Cohen’s d = -0.389), while the Navigation group showed no significant change (t(26) = 0.381, p = 0.706, Cohen’s d = 0.08). When controlling for sex, and site, the Video Control group continued to show the greatest increase in average number of words recalled from pre-test to post-test (t(21) = 3.362, p = 0.003, Marginal R^2^ = 0.134) compared to the Navigation group (t(27) = 0.388, p = 0.701, Marginal R^2^ = 0.021) and the Verbal Memory group (t(26) = -2.307, p = 0.029, Marginal R^2^ =0.083).

A mixed-design ANOVA was conducted with the number of trials to criterion as the dependent variable, condition (Navigation/Verbal Memory/Video Control) as a between-subjects factor, and session (pre/post) as a within-subjects factor. The analysis revealed a significant main effect of Session (F(1, 71) = 25.157, p < 0.001, η^2^ = .0.071) and a marginally significant interaction between Session and Condition (F(2,71) = 2.548, p =0.085, η^2^ = .0.014). Paired-sample t-tests indicated that both the Video Control group (t(20) = -3.65, p = 0.002, Cohen’s d = 0.856) and the Navigation group (t(26) = 3.36, p = 0.002, Cohen’s d = 0.579) demonstrated significant decreases in the number of trials from pre-test to post-test. In contrast, the Verbal Memory group did not show a significant decrease (t(25) = -1.43, p = 0.166, Cohen’s d = 0.238). When controlling for sex and site, the Video Control group (t(21) = -3.74, p = 0.001, Marginal R^2^ = 0.261) and the Navigation group (t(27) = -3.422, p = 0.002, Marginal R^2^ = 0.207) continued to exhibit significant decreases in learning rate from pre to post-test, while the Verbal Memory group (t(26) = -1.456, p = 0.157, Marginal R^2^ = 0.070) did not show a significant change.

***Navigation Transfer task***

*Control analysis for Learning rate of Navigation Transfer task*

When controlling for sex and site using linear mixed effect models or linear regression models, the Navigation group still showed the increase in learning rate from pre to post (t(50) = 4.766, p < 0.001, partial Eta^2^ = 0.312; Verbal Memory group (t(50) = 2.265, p = 0.028, partial Eta^2^ = 0.093; Video Control group (t(19) = 4.081, p < 0.001, marginal R^2^ = 0.358).

*Path error of Navigation Transfer task*

A mixed-design ANOVA was conducted with path error as the dependent variable, condition (Navigation/Verbal Memory/Video Control) as a between-subjects factor, and session (pre/post) as a within-subjects factor. The analysis revealed a significant main effect of session (F(1,70) = 144.97, p < 0.001, η² = 0.443), indicating a substantial change across sessions. However, there was no significant main effect of condition (F(2,70) = 0.726, p = 0.487, η² = 0.007) and no significant session × condition interaction (F(2,70) = 0.59, p = 0.557, η² = 0.004). Paired-sample t-tests indicated that all three groups demonstrated an increase in overall pointing error from post-test to pre-test, with the Navigation group showing the largest effect (Navigation: t(26) = 8.15, p < 0.001, Cohen’s d = 1.97; Verbal Memory: t(26) = 6.47, p < 0.001, Cohen’s d = 1.48; Video Control: t(18) = 6.33, p < 0.001, Cohen’s d = 1.25). When controlling for sex, and site, the Navigation group still showed the greatest increase in overall pointing error from pre-test to post-test (t(26) = 8.307, p < 0.001, Marginal R^2^ = 0.582) compared to Verbal Memory group (t(26) = 6.595, p < 0.001, Marginal R^2^ = 0.416) and Video Control group (t(18) = 6.506, p < 0.001, Marginal R^2^ = 0.517).

During the pre-test session and post-test session, following the navigation task in the virtual Silcton, participants also completed two additional related tasks: the Navigation Pointing task and the Navigation Model Building task (see Methods).

*Navigation Pointing task performance:*

For the pointing task, the error for each trial was determined by calculating the absolute angular difference between the participant's response and the correct angle. These differences were then averaged across all trials to produce an overall pointing error score. We also examined performance differences between participants based on the two types of pointing trials: within-environment and between-environment. We separated trials based on whether the target building was in the environment that the participant was currently standing in (within-environment) or in the other environment (between-environment).

A mixed-design ANOVA was conducted with overall pointing error as the dependent variable, condition (Navigation/Verbal Memory/Video Control) as a between-subjects factor, and session (pre/post) as a within-subjects factor. The analysis revealed a significant main effect of session (F(1,70) = 81.66, p < 0.001, η² = 0.165), but no significant main effect of condition (F(2,70) = 2.618, p = 0.080, η² = 0.048) and no significant interaction between condition and session (F(2,70) = 0.63, p = 0.54, η² = 0.003). Paired-sample t-tests indicated that all three groups demonstrated a decrease in overall pointing error from pre-test to post-test, with the Verbal Memory group showing the largest effect (Navigation: t(26) = 4.76, p < 0.001, Cohen’s d = 0.925; Verbal Memory: t(24) = 5.97, p < 0.001, Cohen’s d = 0.983; Video Control: t(20) = 5.55, p < 0.001, Cohen’s d = 0.758, Supplementary Figure 2a). When controlling for sex, and site in linear mixed effect models, both the Navigation (t(26) = 4.849, p < 0.001, Marginal R^2^ = 0.32) and Verbal Memory group (t(26) = 6.091, p < 0.001, Marginal R^2^ = 0.322) showed the greatest decrease in overall pointing error from pre-test to post-test compared to Video Control group (t(20) = 5.684, p < 0.001, Marginal R^2^ = 0.183).

A mixed-design ANOVA was conducted with within-environment pointing error as the dependent variable, condition (Navigation/Verbal Memory/Video Control) as a between-subjects factor, and session (pre/post) as a within-subjects factor. The analysis revealed a significant main effect of session (F(1,70) = 27.58, p < 0.001, η² = 0.059), but no significant interaction between condition and session (F(2,70) = 0.182, p = 0.834, , η² < 0.001). Paired-sample t-tests indicated that all three groups demonstrated an decrease in within-environment pointing error from pre to post (Navigation: t(26) = 2.36, p = 0.03, Cohen’s d = 0.479; Verbal Memory: t(24) = 3.74, p = 0.001, Cohen’s d = 0.5; Video Control: t(20) = 3.34, p = 0.003, Cohen’s d = 0.435; Supplementary Figure 2c). When controlling for sex and site, all three groups showed a significant decrease in within-environment pointing error from pre to post-test (Navigation: t(26) = 2.403, p = 0.023, Marginal R^2^ = 0.165; Verbal Memory group (t(24) = 3.819, p < 0.001, Marginal R^2^ = 0.213; Video Control group (t(20) = 3.418, p = 0.003, Marginal R^2^ = 0.168).

A mixed-design ANOVA was conducted with between-environment pointing error as the dependent variable, condition (Navigation/Verbal Memory/Video Control) as a between-subjects factor, and session (pre/post) as a within-subjects factor. The analysis revealed a significant main effect of session (F(1,70) = 76.11, p < 0.001, η² = 0.201), but no significant interaction between condition and session (F(2,70) = 0.97, p = 0.39, , η² = 0.005). Paired-sample t-tests indicated that all three groups demonstrated an decrease in between-environment pointing error from pre to post-test (Navigation: t(26) = 5.16, p < 0.001, Cohen’s d = 1.07; Verbal Memory: t(24) = 5.88, p < 0.001, Cohen’s d = 1.13; Video Control: t(20) = 4.22, p < 0.001, Cohen’s d = 0.871; Supplementary Figure 2b). When controlling for sex and site, all three groups showed a significant increase in between-environment pointing error from pre-test to post-test with Navigation group showing the greatest effect (Navigation: t(26) = 5.256, p < 0.001, = 1.17, Marginal R^2^ = 0.360 Verbal Memory group (t(24) = 6.00, p < 0.001, Marginal R^2^ = 0.333; Video Control group (t(20) = 4.320, p < 0.001, Cohen’s d = 0.183).

*Navigation Model Building task performance:*

Map accuracy on the model-building task was measured using a bidimensional regression analysis. A mixed-design ANOVA, with condition (Navigation/Verbal Memory/Video Control) as a between-subjects factor and Session (pre/post) as a within-subjects factor, revealed a significant main effect of session (F(1,72) = 58.799, p < 0.001, η² = 0.144) with no significant interaction between Condition and Session (F(2,72) = 0.957, p = 0.389, η² = 0.005). Paired-sample t-tests indicated that all three groups demonstrated an improvement in map accuracy from pre to post-test (Navigation: t(26) = 4.95, p < 0.001, Cohen’s d = 0.81; Verbal Memory: t(26) = 5.72, p < 0.001, Cohen’s d = 0.94; Video Control: t(20) = 2.97, p = 0.008, Cohen’s d = 0.66; Supplementary Figure 2d). When controlling for sex and site, all three groups still showed the increase in map accuracy from pre-test to post-test (Navigation: t(26) = 5.042, p < 0.001, Marginal R^2^ = 0.272; Verbal Memory: t(26) = 5.825, p < 0.001, Marginal R^2^ = 0.371; Video Control group: t(20) = 3.039, p = 0.006, Marginal R^2^ = 0.119).

***Source memory task performance***

A series of mixed-design ANOVAs was conducted to analyze the effects of condition (Navigation/Verbal Memory/Control; between-subjects factor) and session (Pre/Post; within-subjects factor) on source memory measures: Hit rate, false alarm (FA), and reaction time (RT).

*Hit Rate for Source Memory:* The ANOVA revealed a significant main effect of session (F(1, 63) = 6.382, p = 0.014, η² = 0.017), but no interaction between session and condition (F(2, 63) = 0.206, p = 0.815, η² = 0.001). Paired-sample t-tests indicated that none of the three conditions demonstrated a significant increase in hit rate from pre to post-test (Supplementary Figure 4a): Verbal Memory: t(23) = 1.429, p = 0.166, Cohen’s d = 0.223; Navigation: t(21) = 1.847, p = 0.079, Cohen’s d = 0.316; Control: t(19) = 1.082, p = 0.293, Cohen’s d = 0.208. These effects remained non-significant after controlling for sex and site using linear mixed effect models (ts <1.891, ps > 0.216).

*FA for Source Memory*: The ANOVA showed no significant main effect of session (F(1, 63) = 0.604, p = 0.440, η² = 0.005) or interaction between session and condition (F(2, 63) = 1.445, p = 0.244, η² = 0.022). Paired-sample t-tests showed no significant changes in FA across sessions for any condition: Verbal Memory: t(23) = 1.477, p = 0.153, Cohen’s d = 0.451; Navigation: t(21) = -0.699, p = 0.492, Cohen’s d = -0.168; Control: t(19) = -0.0005, p = 0.999, Cohen’s d = -0.0001, Supplementary Figure 4b. These effects remained non-significant after controlling for sex and site using linear mixed-effect models (ts < 1.575, ps > 0.122).

*Hit Rate for Spatial Source Memory:* The ANOVA revealed a significant main effect of session (F(1, 63) = 5.824, p = 0.019, η² = 0.016), with no interaction between session and condition (F(2, 63) = 0.137, p = 0.872, η² < 0.001). Paired-sample t-tests indicated no significant increase in hit rate from pre-test to post-test in any condition: Verbal Memory: t(23) = 1.102, p = 0.282, Cohen’s d = 0.217; Navigation: t(21) = 1.690, p = 0.106, Cohen’s d = 0.304; Control: t(19) = 1.433, p = 0.168, Cohen’s d = 0.214, Supplementary Figure 4c. These effects remained non-significant after controlling for sex and site using linear mixed-effect models (ts < 1.730, ps > 0.098).

*Hit Rate for Temporal Source Memory:* The ANOVA showed a marginal main effect of session (F(1, 63) = 3.577, p = 0.063. η² = 0.012), with no interaction between session and condition (F(2, 63) = 0.199, p = 0.820, η² = 0.001). Paired-sample t-tests revealed no significant changes in hit rate across sessions: Verbal Memory: t(23) = 1.288, p = 0.210, Cohen’s d = 0.196; Navigation: t(21) = 1.513, p = 0.145, Cohen’s d = 0.288; Control: t(19) = 0.564, p = 0.580, Cohen’s d = 0.147, Supplementary Figure 4d. These effects remained non-significant after controlling for sex and site using linear mixed-effect models (ts < 1.548, ps > 0.136).

*RT for Source Memory:* Although no training effects were observed for hit rate or FA, RT in the source memory task was further examined. A mixed-design ANOVA revealed a marginal main effect of session (F(1, 63) = 3.685, p = 0.059, η² = 0.017) but no significant interaction between session and condition (F(2, 63) = 1.642, p = 0.202, η² = 0.015). Paired-sample t-tests showed no significant changes in RT from pre-test to post-test for the Navigation condition (t(21) = -0.453, p = 0.655, Cohen’s d = -0.107) or the Video Control condition (t(19) = 1.454, p = 0.162, Cohen’s d = 0.363). However, the Verbal Memory condition demonstrated a marginally significant decrease in RT from pre to post-test (t(23) = 2.042, p = 0.053, uncorrected, Cohen’s d = 0.447, Supplementary Figure 5a). This effect remained the same after controlling sex and site using linear mixed-effect model (Verbal Memory: t(24) = 2.086, p = 0.048, Marginal R^2^ = 0.172; Navigation: t(22) = 0.464, p = 0.647, Marginal R^2^ = 0.028; Video Control: t(20) = 1.492, p = 0.151, Marginal R^2^ = 0.111).

Subsequent analyses of spatial and temporal source memory RT revealed a nuanced pattern. Specifically, the Verbal Memory group demonstrated a trend toward decreased RT in temporal source memory from pre to post-test (t(23) = 1.910, p = 0.069, uncorrected, Cohen's d = 0.434, Supplementary Figure 5b), while no such trends were observed in the Navigation (t(21) = -0.622, p = 0.541, Cohen's d = -0.163) or Video Control conditions (t(19) = 1.708, p = 0.103, Cohen's d = 0.445). This effect remained the same after controlling sex and site using linear mixed effect model (Verbal Memory: t(24) = 1.951, p = 0.063, Marginal R^2^ = 0.129; Navigation: t(22) = 0.636, p = 0.531, Marginal R^2^ = 0.038; Video Control: t(20) = 1.752, p = 0.095, Marginal R^2^ = 0.128). Conversely, for spatial source memory, paired-sample t-tests showed no statistically significant RT changes across conditions: Verbal Memory (t(23) = 1.444, p = 0.162, Cohen's d = 0.321), Navigation (t(21) = -0.189, p = 0.852, Cohen's d = -0.041), and Video Control (t(19) = 1.073, p = 0.297, Cohen's d = 0.264. Supplementary Figure 5c).These effects remained not significant after controlling for sex and site using linear mixed-effect models (ts < 1.475, ps > 0.153).

**Supplementary Note** **4:** **Diffusion MRI metrics did not change due to the training**

For DWI, we separately evaluated FW and fwcFA as these measures capture different aspects of tissue microstructure. We also chose to analyze ROIs in gray matter (GM) and white matter (WM) separately, as these tissue types have dramatically different values for both FW and fwcFA. There were 8 GM ROIs and 7 WM ROIs (see Methods). For each metric and tissue type, we performed a mixed-design ANOVA with condition (Navigation/Verbal Memory/Video Control) as a between-subjects factor, ROI and session (pre/post) as within-subjects factors.

For the GM FW analysis, we found no main effect of session (F(1,57) = 0.039, p = 0.844, η^2^ < 0.001, BF_10_ = 0.067, strong evidence against the inclusion of the main effect of session). As expected, we observed a significant main effect of ROI (F(4.397,250.643) = 436.227, p < 0.001, Greenhouse-Geisser corrected, η^2^ = 0.933, BF_10_ > 100, extremely strong evidence for the inclusion of the main effect of ROI) and a significant main effect of condition (F(2,57) = 3.418, p = 0.040, η^2^ = 0.007, BF_10_ = 0.879, strong evidence against including the main effect of condition). However, we found no significant interaction between session × condition × ROI (F(6.169,175.825) = 0.376, p =0.898, BF_10_ < 0.001, strong evidence against this interaction). Paired-sample t-tests confirmed that none of the three training groups exhibited significant changes from pre to post-test in any of the 8 ROIs (*ps* > 0. 842, FDR corrected). These findings remained nonsignificant after controlling for sex and site as covariates (*ps* > 0.826, FDR corrected).

For the GM fwcFA analysis, we found no main effect of session (F(1,57) = 0.471, p = 0.495, η^2^ < 0.001, BF_10_ = 0.090, strong evidence against the inclusion of the main effect of session). As expected, we observed a significant main effect of ROI (F(3.584,204.312) = 207.954, p < 0.001, Greenhouse-Geisser corrected, η^2^ = 0.637, BF_10_ > 100, extremely strong evidence for the inclusion of the main effect of ROI). There was no significant main effect of condition (F(2,57) = 1.147, p = 0.325, η^2^ = 0.005, BF_10_ = 0.011, strong evidence against including the main effect of condition). We found no significant interaction between session × condition × ROI (F(4.928,140.441) = 0.288, p = 0.917, η^2^ < 0.001, BF_10_ < 0.001, strong evidence against this interaction). Paired-sample t-tests confirmed that none of the three training groups exhibited significant changes from pre to post-test in any of the 8 ROIs (*ps* > 0. 958, FDR corrected). These findings remained not significant after controlling for sex and site as covariates (*ps* > 0.834, FDR corrected).

For the WM FW analysis, we found no main effect of session (F(1,57) = 0.012, p = 0.913, η^2^ < 0.001, BF_10_ = 0.767, strong evidence against the inclusion of the main effect of session). As expected, we observed a significant main effect of ROI (F(2.938, 167.445) = 1659.245, p < 0.001, Greenhouse-Geisser corrected, η^2^ = 0.791, BF_10_ > 100, extremely strong evidence for the inclusion of the main effect of ROI). There was no significant main effect of condition (F(2,57) = 0.1, p = 0.905, η^2^ < 0.001, BF_10_ = 0.431, strong evidence against including the main effect of condition). We found no significant interaction between session × condition × ROI (F(7.623, 217.255) = 0.923, p = 0.495, η^2^ < 0.001, BF_10_ < 0.001, strong evidence against this interaction). Paired-sample t-tests confirmed that none of the three training groups exhibited significant changes from pre to post-test in any of the 7 ROIs (*ps* > 0. 400, FDR corrected). These findings remained not significant after controlling for sex and site as covariates (*ps* > 0.597, FDR corrected).

For the WM fwcFA analysis, we found no main effect of session (F(1,57) < 0.001, p = 0.981, η^2^ < 0.001, BF_10_ = 0.090, strong evidence against the inclusion of the main effect of session). As expected, we observed a significant main effect of ROI (F(4.093, 233.279) = 2340.878, p < 0.001, Greenhouse-Geisser corrected, η^2^ = 0.628, BF_10_ > 100, extremely strong evidence for the inclusion of the main effect of ROI). There was no significant main effect of condition (F(2,57) = 0.025, p = 0.975, η^2^ < 0.001, BF_10_ = 0.011, strong evidence against including the main effect of condition). We found no significant interaction between session × condition × ROI (F(6.924, 197.328) = 1.954, p = 0.064, η^2^ < 0.001, BF_10_ < 0.001, strong evidence against this interaction). Paired-sample t-tests confirmed that none of the three training groups exhibited significant changes from pre to post-test in any of the 7 ROIs (*ps* > 0. 083, FDR corrected). These findings remained not significant after controlling for sex and site as covariates (*ps* > 0.063, FDR corrected).

**Supplementary Note 5: The correlation between hippocampal volume and behavioral performance**

**Improvements in the learning rate of the Verbal Memory Transfer task, but not the Navigation Transfer task, were found to correlate with lateral hippocampal volume but not with anterior and posterior hippocampus (using average volume between pre and post-test).**

Consistent findings were obtained when analyzing hemispheric hippocampal volumes (see Supplemental Figure 3). Specifically, a significant positive correlation was observed between the learning rate in the Verbal Memory group and both left hippocampal volume (r(23) = 0.568, p-FDR = .014) and right hippocampal volume (r(23) = 0.601, p-FDR = .007), while no significant correlations were found for the Navigation (left: r(25) = 0.038, p-FDR = .858; right: r(25) = 0.008, p-FDR = .970) or Video Control group (left: r(19) = -0.123, p-FDR = .858; right: r(19) = -0.163, p = .758), after controlling for sex and site as covariates. Fisher's z-tests revealed that the positive correlation in the Verbal Memory group was significantly stronger than that in the Navigation group even after controlling for sex and site (left hippocampus: Z = 1.942, p-FDR = 0.039; right hippocampus: Z = 2.326, p-FDR = 0.015) and Video Control group (left hippocampus: Z = 2.234, p-FDR = 0.038; right hippocampus: Z = 2.702, p-FDR = 0.010) while no significant difference was observed between the Navigation group and Video Control group (left hippocampus: Z = -0.439, p = 0.669; right hippocampus: Z = -0.553, p-FDR = 0.710).

An analysis was conducted to assess the correlation between anterior and posterior hippocampal volumes and changes in learning rate on the verbal memory transfer task. However, the improvement in learning rate demonstrated no significant correlation with either anterior or posterior hippocampal volume in any of the three groups (ps > 0.232, Supplementary Table 4). This lack of correlation remained consistent after accounting for the influence of sex and site as covariates (ps > 0.114).

We also correlated hippocampal volume and MTL subregions with the change in the average number of words recalled, number of trials to criterion, and slope from linear regression (see Supplementary Note 3) between the post-test and pre-test for the verbal memory transfer task. The analysis revealed no significant correlations between hippocampal volume or MTL subregions and either the average number of words recalled or the number of trials to criterion (Supplementary Tables 6 and 7), regardless of whether sex and site were included as covariates. However, when correlating slope with hippocampal volume, a positive correlation was found for the Verbal Memory group (total hippocampal volume: r = 0.552, p = 0.006; left hippocampus: r = 0.502, p = 0.015 ;right hippocampus: r = 0.561, p = 0.005, Supplementary Table 8), however, no such correlation was found for the Navigation group (total hippocampus: r = 0.094, p = 0.656; left hippocampus: r = 0.114, p = 0.586; right hippocampus: r = 0.073, p = 0.727) and Video Control (total hippocampus: r = -0.239, p = 0.325; left hippocampus: r = -0.217, p = 0.372; right hippocampus: r = -0.238, p = 0.326) even after controlling for sex and site as covariates.

We also correlated hippocampal volume with the change in learning rate from the post-test and pre-test for the navigation transfer task. No significant correlations were found between total hippocampal volume and changes in learning rate in any of the three conditions (Navigation: r(25) = -0.27, p = .181; Verbal Memory: r(24) = -0.31, p = .129; Video Control: r(17) = 0.124, p = .612), regardless of whether sex and site were included as covariates. Similar no significant correlations were observed for both left hippocampal volume (Navigation: r(25) = -0.25, p = .214; Verbal Memory: r(24) = -0.29, p = .160; Video Control: r(17) = 0.25, p = .310) and right hippocampal volume (Navigation: r(25) = -0.27, p = .167; Verbal Memory: r(24) = -0.302, p = .142; Video Control: r(17) = -0.018, p = .940), regardless of whether sex and site were included as covariates. We also did not find any significant correlations when examining hippocampal subfields or surrounding MTL subregions (p’s > 0.20, Supplementary Table 5).

The relationship between total hippocampal volume, MTL subregion volumes and changes in performance on the navigation transfer task was assessed through correlational analyses, focusing on path error, overall pointing error, between-environment pointing error, within-environment pointing error, and map accuracy. These analyses demonstrated a lack of significant associations between hippocampal volume and MTL subregions and any of the behavioral metrics under consideration (Supplemental Tables 9-13), regardless of the inclusion of sex and site as covariates.

**Improvements in the learning rate of the Verbal Memory Transfer task, but not the Navigation Transfer task, were found to correlate with both total hippocampal volume and the volume of the CA2/3/DG subfield (using volume data only from the pre-test).** We examined the hypothesis that baseline hippocampal volume might be associated with either verbal or navigation performance. For this analysis, we utilized the hippocampal volume (or subfield volume) obtained for each participant at pre-test, rather than averaging volumes across pre and post-test. We then assessed the relationship between hippocampal volumes at pre-test and the change in learning rate from pre-test to post-test for both the Verbal Memory transfer task and the Navigation Transfer task.

We found a marginal correlation in the verbal memory training group between the pre-test total hippocampal volume and the observed improvement in verbal memory performance from pre- to post-test (r(23) = 0.352, p = .084). Further analysis accounting for sex and site as covariates revealed a significant positive correlation between total hippocampal volume and the learning rate in the Verbal Memory condition (r(23) = 0.545, p = .007). This effect was specific to the verbal memory training; no significant correlation was identified for either the Navigation condition (r(25) = 0.072, p = .721) or the Video condition (r(19) = -0.209, p = .362); the same result was observed when controlling for covariates (Navigation condition: r(25) = 0.082, p = .695; Video condition: r(19) = -0.144, p = .555). Consistent findings were obtained when analyzing hemispheric hippocampal volumes. Specifically, a significant positive correlation was observed between the learning rate in the Verbal Memory condition and both left hippocampal volume (r(23) = 0.504, p = .014) and right hippocampal volume (r(23) = 0.545, p = .007), while no significant correlations were found for the Navigation condition (left: r(25) = 0.122, p = .561; right: r(25) = 0.045, p = .832) or Video Control condition (left: r(19) = -0.084, p = .734; right: r(19) = -0.187, p = .444), after controlling for sex and site as covariates.

We further examined the correlation between CA23DG volume and learning rate change. The CA23DG subfield showed a positive correlation with the change in learning rate from pre to post-test in the verbal memory transfer task of the Verbal Memory condition (r(23) = 0.504, p = .01), suggesting that individuals in the Verbal Memory condition with larger CA23DG volumes exhibited greater improvements in memory performance from pre- to post-test. This correlation persisted even after controlling for sex and site as covariates (r(23) = 0.439, p = .036). No significant correlations were observed for the CA23DG subfield in the Navigation (r(25) = 0.007, p = .972) or Video Control conditions (r(19) = -0.05, p = .828), regardless of whether sex and site were included as covariates.

Fisher's z-tests revealed that the positive correlation in the Verbal Memory condition was significantly stronger than that in the Navigation even after controlling for sex and site (total hippocampus: Z = 1.793, p = 0.037; left hippocampus: Z= 1.464, p = 0.072; right hippocampus: Z= 1.919, p = 0.027; CA23DG: Z = 1.687, p = .046) and Video conditions (total hippocampus: Z = 2.382, p = 0.009; left hippocampus: Z = 2.009, p = 0.022; right hippocampus: Z= 2.518, p = 0.006; CA23DG: Z = 1.766, p = .038), while no significant difference was observed between the Navigation and Video Control conditions (total hippocampus: Z = -0.731, p = 0.768; left hippocampus: Z= -0.662, p = 0.746; right hippocampus: Z= -0.750, p = 0.773; CA23DG: Z = -0.214, p = 0.585).

**Improvements in the learning rate of the Verbal Memory Transfer task, but not the Navigation Transfer task, were found to correlate with both total hippocampal volume and the volume of the CA2/3/DG subfield (using volume data only from the post-test).** We examined the hypothesis that baseline hippocampal volume may be associated with either verbal or navigation performance. For this analysis, we utilized the hippocampal volume (or subfield volume) obtained for each participant at post-test, rather than averaging volumes across pre-test and post-test. We then assessed the relationship between hippocampal volumes at post-test and the change in learning rate from pre-test to post-test for both the Verbal Memory transfer task and the Navigation transfer task.

We found a marginal correlation in the verbal memory training group between the post-test total hippocampal volume and the observed improvement in verbal memory performance from pre- to post-test (r(23) = 0.360, p = .078). Further analysis accounting for sex and site as covariates revealed a significant positive correlation between total hippocampal volume and the learning rate in the Verbal Memory condition (r(23) = 0.623, p = .001). This effect was specific to the verbal memory training; no significant correlation was identified for either the Navigation condition (r(25) = -0.009, p = .965) or the Video Control condition (r(19) = -0.178, p = .439); the same was true when controlling for covariates (Navigation condition: r(25) = 0.000, p = .999; Video Control condition r(18) = -0.083, p = .736). Consistent findings were obtained when analyzing hippocampal volumes by hemisphere. Specifically, a significant positive correlation was observed between the learning rate in the Verbal Memory condition and both left hippocampal volume (r(23) = 0.597, p = .003) and right hippocampal volume (r(23) = 0.594, p = .003), while no significant correlations were found for the Navigation (left: r(25) = 0.026, p = .901; right: r(24) = -0.023, p = .912) or Video Control conditions (left: r(19) = -0.036, p = .883; right: r(18) = -0.109, p = .656), after controlling for sex and site as covariates.

We further examined the correlation between CA23DG volume and learning rate change. The CA23DG subfield showed a positive correlation with the change in learning rate from pre to post-test in the verbal memory transfer task of the Verbal Memory condition (r(23) = 0.545, p = .005), suggesting that individuals in the Verbal Memory condition with larger CA23DG volumes exhibited greater improvement in memory performance from pre to post-test. This correlation persisted even after controlling for sex and site as covariates (r(23) = 0.537, p = .008). No significant correlations were observed for the CA23DG subfield in the Navigation (r(25) = -0.027, p = .894) or Video Control conditions (r(19) = -0.110, p = .634), regardless of whether sex and site were included as covariates.

Fisher's z-tests revealed that the positive correlation in the Verbal Memory condition was significantly stronger than that in the Navigation group even after controlling for sex and site (total hippocampus: Z = 2.473, p = 0.007; left hippocampus: Z = 2.243, p = 0.012; right hippocampus: Z = 2.398, p = 0.008; CA23DG: Z = 2.209, p = .014) and the Video Control condition (total hippocampus: Z = 2.558, p = 0.005; left hippocampus: Z = 2.280, p = 0.011; right hippocampus: Z = 2.498, p = 0.006; CA23DG: Z = 2.337, p = .014), while no significant difference was observed between the Navigation and the Video conditions (total hippocampus: Z = -0.266, p = 0.605; left hippocampus: Z = -0.2, p = 0.841; right hippocampus: Z = -0.276, p = 0.609; CA23DG: Z = -0.291, p = 0.615).

***The correlation between the Verbal Memory Training task, hippocampus volume and MTL subfields***

Additionally, we calculated correlations between the slopes from the first five days and the last five days of the Verbal Memory training task with hippocampal volume and MTL subregions separately. No significant correlations were identified between the training effect and any MTL subregion or the total hippocampal volume (Supplemental Tables 14 and 15).

**Supplementary Note 6: Informational connectivity results**

***Informational connectivity changes specific to spatial or temporal context encoding in the Verbal Memory and Navigation interventions***

In the Verbal Memory condition, increased informational connectivity during encoding was observed during the post-test stage compared to the pre-test stage (i.e., post > pre), relative to the combined Navigation and Video conditions. Specifically, enhanced connectivity was identified between the right frontal orbital cortex and right precuneus (t = 3.93, p < 0.05), the right dorsal lateral occipital cortex (dLOC) and left temporal occipital fusiform cortex (TOFC) (t = 3.98, p < 0.05), and the right temporal pole and left superior parietal lobule (SPL) (t = 4.33, p < 0.05). Conversely, no significant decreases in connectivity were observed from post to pre-test (i.e., post < pre) in the Verbal Memory condition, relative to the combined Navigation and Video conditions (Figure 4d, top-middle). In contrast, the Navigation condition showed significantly decreased informational connectivity during the post-test compared to the pre-test stage (i.e., post < pre), relative to the combined Verbal Memory and Video conditions. Specifically, reduced connectivity was observed between the right intracalcarine cortex and the left ventral LOC (t = 3.85, p < 0.05). However, the Navigation condition also demonstrated significantly increased connectivity during post compared to pre (i.e., post > pre) between the left posterior cingulate gyrus (PCC) and right temporal fusiform cortex (TFC) (t = 4.44, p < 0.05), as well as between the left middle temporal gyrus and left SPL (t = 4.96, p < 0.05), relative to the combined Verbal Memory and Video conditions (Figure 4d, bottom-middle).

Similar patterns emerged when analyzing trials encoded within temporal contexts. In the Verbal Memory condition, increased informational connectivity was observed during the post-test compared to the pre-test (i.e., post > pre), relative to the combined Navigation and Video conditions. Specifically, enhanced connectivity was identified between the left middle frontal gyrus (MFG) and left middle temporal gyrus (MTG) (t = 4.01, p < 0.05). Conversely, no significant decreases in connectivity were observed from post to pre (i.e., post < pre) in the Verbal Memory condition, relative to the combined Navigation and Video conditions (Figure 4d, top-right). In contrast, the Navigation condition exhibited significantly decreased informational connectivity during the post-test compared to the pre-test (i.e., post < pre), relative to the combined Verbal Memory and Video conditions. Specifically, reductions were observed between the left frontal orbital cortex and right angular gyrus (AG) (t = 4.07, p < 0.05), right supramarginal gyrus (RSMG) and left supramarginal gyrus (LSMG) (t = 3.63, p < 0.05), left superior frontal gyrus (LSFG) and right frontal pole (t = 4.18, p < 0.05), left paracingulate gyrus and right frontal pole (t = 4.04, p < 0.05), and LSFG and right frontal pole (t = 4.14, p < 0.05). Conversely, no significant increases in connectivity were observed from post to pre (i.e., post > pre) in the Navigation condition, relative to the combined Verbal Memory and Video conditions (Figure 4d, bottom-right).

***Informational connectivity changes specific to spatial or temporal context retrieval in the Verbal Memory and Navigation interventions***

When analyzing only spatial RSMs during source retrieval, we observed decreased informational connectivity between the left superior frontal gyrus (LSFG) and right superior frontal gyrus (SFG) (t = 4.50, p < 0.05), as well as between the right frontal pole and right posterior cingulate cortex (PCC) (t = 4.10, p < 0.05) in the post-test compared to the pre-test (i.e., post < pre) in the Verbal Memory condition, relative to the combined Navigation and Video conditions. Conversely, we observed increased connectivity between the left middle temporal gyrus (LMTG) and right supramarginal gyrus (RSMG) (t = 3.51, p < 0.05), and between the right dorsal lateral occipital cortex (RdLOC) and right supramarginal gyrus (RSMG) (t = 3.73, p < 0.05) from pre to post (i.e., post > pre) in the Verbal Memory condition, relative to the combined Navigation and Video conditions (Figure 5b, top-middle).

We observed significantly increased informational connectivity between the left frontal pole and right dorsal lateral occipital cortex (dLOC) (t = 3.42, p < 0.05), as well as between the left frontal pole and right parahippocampal cortex (PHC) (t = 4.32, p < 0.05) in the post-test compared to the pre-test (i.e., post > pre) in the Navigation condition, relative to the combined Verbal Memory and Video conditions. Conversely, no significant decreases in connectivity were observed from pre to post (i.e., post < pre) in the Navigation condition, relative to the combined Verbal Memory and Video conditions (Figure 5b, bottom-middle).

When analyzing only temporal RSMs during source retrieval, we observed significantly decreased informational connectivity from pre-test to post-test (i.e., post < pre) in the Verbal Memory condition, relative to the combined Navigation and Video conditions, across multiple brain regions. Specifically, decreased connectivity was found between the following regions: left frontal pole and left middle temporal gyrus (t = 3.71, p < 0.05), left frontal pole and left angular gyrus (t = 4.03, p < 0.05), left paracingulate gyrus and left precuneus (t = 3.62, p < 0.05), left precuneus and left ventral lateral occipital cortex (t = 3.60, p < 0.05), left ventral lateral occipital cortex and right dorsal lateral occipital cortex (t = 4.12, p < 0.05), left frontal pole and right angular gyrus (t = 4.70, p < 0.05), left paracingulate gyrus and right angular gyrus (t = 4.39, p < 0.05), left paracingulate gyrus and right middle temporal gyrus (t = 4.12, p < 0.05), right precuneus and right frontal pole (t = 4.68, p < 0.05), left medial frontal gyrus and right inferior frontal gyrus (t = 3.91, p < 0.05), left precuneus and right middle frontal gyrus (t = 3.91, p < 0.05), left precuneus and right dorsal lateral occipital cortex (t = 3.49, p < 0.05), right paracingulate gyrus and right dorsal lateral occipital cortex (t = 4.06, p < 0.05), left retrosplenial cortex and right medial frontal cortex (t = 3.99, p < 0.05), left frontal pole and right frontal pole (t = 4.81, p < 0.05), left frontal pole and right superior frontal gyrus (t = 3.90, p < 0.05), and left frontal pole and right precuneus (t = 4.14, p < 0.05), as well as right precuneus and left superior frontal gyrus (t = 4.14, p < 0.05). In contrast, we observed an increase in connectivity between the left ventral lateral occipital cortex and left middle frontal gyrus (t = 3.96, p < 0.05) from pre to post (i.e., post > pre) in the Free Recall condition, relative to the combined Navigation and Video conditions (Figure 5b, top-right).

We observed a significant increase in informational connectivity from pre-test to post-test (i.e., post > pre)in the Navigation condition, relative to the combined Verbal Memory and Video conditions, between the following regions: left precuneus and left middle frontal gyrus (t = 4.10, p < 0.05), left precuneus and left superior frontal gyrus (t = 3.89, p < 0.05), right lingual gyrus and right dorsal lateral occipital cortex (t = 4.07, p < 0.05), right lingual gyrus and right dorsal lateral occipital cortex (t = 3.68, p < 0.05), and left superior frontal gyrus and right middle temporal gyrus (t = 3.77, p < 0.05). In contrast, no significant decreases in connectivity were observed from pre to post (i.e., post < pre) in the Navigation condition, relative to the combined Verbal Memory and Video conditions (Figure 5b, bottom-right).

**Supplementary Note 7: Univariate activation results**

***Task-related changes in brain activation during encoding as a result of both the Verbal Memory and Navigation interventions***

We investigated how different training interventions influenced univariate brain activation during the encoding phase by examining whole-brain activity changes between the pre-test and post-test stages for the Verbal Memory, Navigation, and Video conditions. In the Verbal Memory condition, a significant decrease in activation was observed in several regions during the post-test compared to the pre-test (post < pre). These regions included the left dorsolateral occipital cortex (dLOC; Z = 4.66, MNI: -46, -62, 30), the left middle frontal gyrus (MFG; Z = 4.32, MNI: -42, 12, 50), the left frontal pole (Z = 4.20, MNI: -24, 40, 46), the left middle temporal gyrus (MTG; Z = 4.28, MNI: -60, -24, -12), and the left precuneus (Z = 3.80, MNI: -10, -52, 38) (see Supplementary Figure 6a). In contrast, in the Navigation condition, post-test activation was significantly increased in the right middle temporal gyrus (MTG; Z = 4.03, MNI: 42, -54, 6) compared to the pre-test (post > pre) (see Supplementary Figure 6b).

However, no group-specific changes in activity were observed when comparing the pre-test and post-test stages across the three conditions: Verbal Memory, Navigation, and Video Control (Supplementary Table 16).

***Activity changes specific to spatial or temporal context encoding in the Verbal Memory and Navigation training conditions***

We also investigated whether navigation and Verbal Memory training differentially affected spatial versus temporal encoding. In the Verbal Memory condition, a significant decrease in spatial encoding activation was observed during the post-test compared to the pre-test (post < pre, Supplementary Figure 6c). Specifically, reduced activation was detected in the left lateral occipital cortex (LOC; Z = 3.97, MNI: -44, -62, 32) and the left frontal pole (Z = 3.99, MNI: -18, 48, 36). Similarly, in the Verbal Memory group, a decrease in temporal encoding activation was observed in the left LOC (Z = 3.97, MNI: -40, -62, 32) during the post-training stage relative to the pre-test (post < pre, Supplementary Figure 6d). In contrast, no significant differences between pre and post were found in the Navigation condition for either spatial or temporal encoding.

***No task-related changes in brain activation during retrieval as a result of either the verbal memory or navigation interventions***

See Supplementary Table 16

***No activity changes specific to spatial or temporal context retrieval in the Verbal Memory and Navigation training conditions***

See Supplementary Table 16
